# Population genomics of *Pocillopora* corals: insights from RAD-sequencing

**DOI:** 10.1101/2021.05.04.442552

**Authors:** Didier Aurelle, Marine Pratlong, Nicolas Oury, Anne Haguenauer, Pauline Gélin, Hélène Magalon, Mehdi Adjeroud, Pascal Romans, Jeremie Vidal-Dupiol, Michel Claereboudt, Camille Noûs, Lauric Reynes, Eve Toulza, François Bonhomme, Guillaume Mitta, Pierre Pontarotti

## Abstract

Scleractinian corals are of great ecological interest as ecosystem engineer species. Accordingly, there is a wealth of studies on their adaptive abilities facing climate change. Such studies should rely on precise species and population delimitation. Nevertheless species delimitation in corals can be hindered by the lack of adequate genetic markers, by hybridization, and by morphological plasticity. Here we applied RAD sequencing to the study of species delimitation and genetic structure in populations of *Pocillopora* spp. from Oman and French Polynesia with the objectives to test primary species hypotheses based on mitochondrial DNA sequencing, and to study the genetic structure among sampling sites inside species. Regarding the varying levels of missing data observed among samples we tested different filtering strategy. The main genetic differentiation was observed between samples from Oman and French Polynesia, which also corresponded to different mitochondrial lineages and species hypotheses. In Oman, we did not observe any clear differentiation according to the main mitochondrial lineages considered here, nor between sampling sites. In French Polynesia where a single mitochondrial lineage was studied, we did not evidence any differentiation according to sampling sites. These results provide an additional example of the importance of using independent nuclear markers for the study of species delimitation. Our analyses also allowed the identification of clonal lineages in our samples, and to take them into account in our interpretations. We used simulations to study the impact of clonal reproduction on the distribution of statistics of genetic diversity and genetic structure among loci.

## Introduction

Anthozoans, i.e. hexacorals and octocorals, are ecologically key species in various marine ecosystems, from tropical coral reefs to deep coral species. They are the subject of numerous studies on the impact of climate change, as heat waves can lead to bleaching or necrosis events in tropical as well as temperate species (Garrabou et al., 2009; Hughes et al., 2018). Anthozoans are also important models in evolutionary biology, from phylogenetic studies to better understand their origin and long-term evolution (Kayal et al., 2018; Pratlong et al., 2017), to population genetics studies dealing with dispersal, parentage analysis or sex determinism (Ledoux et al., 2010; Mokhtar-Jamaï et al., 2013; Pratlong et al., 2017; Sheets et al., 2018; Underwood et al., 2007). These studies rely on adequate species delineation, as for example to discuss the diversity of thermotolerance or of patterns of connectivity (Brener-Raffali et al., 2019; Pante et al., 2015). Nevertheless in Anthozoans, species limits can be difficult to infer because of morphological plasticity, slow evolution of mitochondrial DNA, or hybridization (Aurelle et al., 2017; Calderon et al., 2006; Gélin et al., 2017; Marti-Puig et al., 2014).

Hexacorals of the *Pocillopora* genus (Lamarck, 1816), such as the morpho-species *P. damicornis, P. eydouxi*, or *P. acuta*, are common corals found in shallow waters in the Red Sea, Indian and Pacific Oceans. The morphological diversity of this genus has led to different taxonomical categorizations. The use of mitochondrial DNA and microsatellite data further led to the proposal of primary and secondary species hypotheses (PSH and SSH), which are not always congruent with species hypotheses based on morphology (i.e. morpho-species: (Gélin et al., 2017). These results raise questions regarding the possibility of hybridization and introgression between different species in the *Pocillopora* genus. The use of partial genomic sequencing, such as Restriction Sites Associated DNA sequencing (RAD-Seq), allows the simultaneous discovery and genotyping of Single Nucleotide Polymorphism (SNPs) in non-model organisms (Baird et al., 2008). Restriction Sites Associated DNA sequencing has thus been used to test species delimitations in octocorals (Pante et al., 2015) and hexacorals (Forsman et al., 2017), including *Pocillopora*. The first RAD-Seq study dealing with *Pocillopora* corals suggested the possibility of hybridization of *P. damicornis* with *P. eydouxi* and *P. elegans* (Combosch & Vollmer, 2015). However, the samples of this last study were pooled for sequencing, which prevented inferences on individual admixture. Through the analysis of additional *Pocillopora* lineages with RAD-Seq, Johnston et al. (2017) found a good concordance with the phylogenetic relationships inferred from mitochondrial DNA data. Their data suggested the possibility of hybridization or incomplete lineage sorting among the closest lineages studied there.

Additional genomic studies of *Pocillopora* spp. populations would be interesting at different levels. First, in situation of sympatry, it would allow testing more precisely the possibility of hybridization (i.e. the existence of individuals with mixed ancestry) or introgression (the resulting gene flow, if any, inside a given gene pool) among putative species. Second, after species delimitation, it would be useful to study the genetic differentiation and connectivity among populations (Pante et al., 2015). Third, inside populations, RAD-Seq could be used to study the potential impact of clonality on the genomic diversity of these corals. Indeed, as in many other scleractinian species, clonality has been demonstrated in *P. acuta* (Pauline Gélin et al., 2017), and this can impact the study of genetic structure (Adjeroud et al., 2014; Balloux et al., 2003).

Here we used a hierarchical sampling design to study the genomic diversity of *Pocillopora* lineages at different scales. Specifically we sampled *Pocillopora* spp. in two distant regions located at the margins of the distribution range of the genus: French Polynesia, Central Pacific Ocean, and Oman, Northwestern Indian Ocean, with different sites in each region. This sampling scheme first aimed at studying the diversity of thermotolerance according to different thermal regimes (see Brener-Raffali et al., 2019). Despite sampling morphologically similar colonies in French Polynesia and Oman, we got samples which corresponded to different mitochondrial haplotypes, and species hypotheses. In Oman several mitochondrial lineages were also sampled in sympatry. We therefore applied RAD-Seq to these samples to test species limits and genetic differentiation among sampling sites. We simulated data with different levels of clonal reproduction to help in the interpretation of results obtained with RAD-Seq.

## Materials and methods

### Sampling and DNA extraction

Five sites were sampled in Oman (hereafter O1 to O5 in sample names, export CITES n° 37/2014 / import CITES n° FR1406600081-I) six sites at two islands in French Polynesia (hereafter Polynesia, with sites MH, MV, MT, at Moorea, and TF, TV; TT at Tahiti; Table 1; export CITES n° FR1398700171-E / import CITES n° FR1306600053-I). Thirty colonies were sampled in each site in Oman (except at O3, with 13 colonies), and ten colonies per site in Polynesia. The sampling was focused on coral colonies morphologically similar with a *P. acuta (damicornis* type *β*)-like *corallum* morphology. Both in Oman and Polynesia, we also sampled additional colonies potentially belonging to other species, to be used as outgroups. The corresponding species hypotheses were checked through the sequencing of part of the mitochondrial DNA (see below).

**Table 1:** characteristics of sampling sites. For the final sampling sizes depending on datasets and mitochondrial lineages, see supplementary table S1. The thermal regime gives a qualitative indication of temperature variability at the corresponding sampling site.

| Region | Site | GPS | Code | Sampling date | Depth (m) | Thermal regime |
| --- | --- | --- | --- | --- | --- | --- |
| Polynesia | Moorea Haapiti | 17°32'39.27 S<br>149°53'37.40 W | MH | 03/2014 | 0.5-2 | Low variations |
| Polynesia | Moorea Tiahura | 17°29'17.41 S<br>149°53'45.58 W | MT | 03/2014 | 0.5-2 | Low variations |
| Polynesia | Moorea Vaiare | 17°31'24.10 S<br>149°46'33.85 W | MV | 03/2014 | 0.5-2 | Low variations |
| Polynesia | Tahiti Faratea | 17°43'17.61 S<br>149°18'11.78 W | TF | 03/2014 | 0.5-2 | Low variations |
| Polynesia | Tahiti Vairao | 17°48'20.90 S<br>149°17'43.13 W | TV | 03/2014 | 0.5-2 | Low variations |
| Polynesia | Tahiti Tautira | 17°45'12.11 S<br>149° 9'26.68 W | TT | 03/2014 | 0.5-2 | Low variations* |
| Oman | Bandar Al Khayral 1 | 23°30'54.25 N<br>58°45'15.70 E | O1 | 06/2014 | 2-8 | High variations |
| Oman | Bandar Al Khayral 21 | 23°31'26.66 N<br>58°44'2.18 E | O2 | 06/2014 | 2-8 | High variations |
| Oman | Bandar Al Khayral 3 | 23°31'8.90 N<br>58°45'29.40 E | O3 | 06/2014 | > 12 | High variations but less than O1, O2 and O4 |
| Oman | Muscat | 23°37'28.61 N<br>58°36'1.39 E | O4 | 06/2014 | 2-8 | High variations |
| Oman | Daymaniat | 23°51'25.12 N<br>58° 6'3.43 E | O5 | 06/2014 | 2-8 | High variations but less than O1, O2 and O4 |

\* see Brener-Raffalli et al. (2019) for further details on thermal regime in Oman.

After sampling, all colonies fragments were bleached with menthol according to previously detailed protocols (Vidal-Dupiol et al., 2019; Wang et al., 2012). Total genomic DNA was extracted according to the protocol of (Sambrook et al., 1989), followed by a purification using Qiagen DNeasy blood and tissue spin columns (Qiagen). Genomic DNA concentration was quantified using a Qubit 2.0 Fluorometer (Life Technologies).

### Mitochondrial ORF sequencing and microsatellite genotyping

We used mitochondrial sequencing and microsatellite genotyping to assign the colonies to Primary Species Hypothesis (PSH) and to Secondary Species Hypothesis (SSH) according to the nomenclature of Gélin et al., (2017) which will be used hereafter. The mitochondrial variable open reading frame (so-called ORF which corresponds to part of the ATP synthase subunit 6 gene) was amplified with the FATP6.1 (5’-TTTGGGSATTCGTTTAGCAG-3’) and RORF (5’- SCCAATATGTTAAACASCATGTCA-3’) primers (Flot & Tillier, 2007) and submitted to Sanger sequencing in both directions. The GenBank accession numbers are indicated in Gélin et al., 2017. Protein-coding sequences were analysed using MEGA version 6 (Tamura et al., 2013). Sequence alignment was performed using MUSCLE, by including mitochondrial sequences used in Gélin et al. (2017). The best model (Kimura-2 parameters with uniform substitution rates) was selected for the lowest BIC (Bayesian Information Criterion). Maximum-likelihood tree was computed with the best model, and the robustness of the tree was tested with 1000 bootstrap replicates. The colonies were then assigned to the corresponding species hypothesis by comparison with the results of Gélin et al. (2017).

Additionally, a subset of colonies (N = 165) were genotyped with 13 microsatellite loci, and assigned to species hypotheses with Bayesian clustering as in Gélin et al. (2017). Furthermore, as PSH05 (*P. acuta* or *P. damicornis* type *β*) is known to propagate asexually (Gélin et al., 2017; Gélin et al., 2018), microsatellites genotyping was used to search for repeated multilocus genotypes (MLGs) as a benchmark for the delimitation of clonal lineages with RAD-Seq.

### RAD sequencing and analyses

The preparation and sequencing of RAD-Seq library was performed as in (Pratlong et al., 2018). We started with an initial number of 211 individuals, distributed among seven libraries. The sequences were first demultiplexed, filtered by quality, and searched for adapters contamination with iPyrad v0.7.28 with default parameters (https://ipyrad.readthedocs.io/). We then checked the quality of the sequences and the absence of adapters with FastQC (Andrews, 2010). The assembly of RAD loci has been done with Stacks 2.3 (Catchen et al., 2013). We used a published genome of *P. damicornis* (Cunning et al., 2018) as a reference to map the reads with BWA (Li & Durbin, 2009). At that step, the levels of reads and of missing data were very uneven among the 211 samples (see results). As a consequence we chose to use different assembling strategies leading to different datasets. Following preliminary assembly analyses, we removed individuals with less than 900 000 reads: their inclusion led to datasets with very low numbers of SNPs. The resulting dataset comprised 140 individuals: 104 from Oman and 36 from Polynesia. Then we used the module populations of stacks to assemble three datasets: 1) one considering all 140 individuals grouped by mitochondrial lineage (“All” dataset), 2) one with only samples from Oman and separation based on mitochondrial lineage, and 3) one with only samples from Polynesia and a separation based on sampling sites (apart from outgroups, all individuals from Polynesia shared the same mitochondrial haplotype; see results). At that stage, the individuals of the All and Oman datasets for which we did not get any mitochondrial sequence were put in an additional group for assembly (“unknown”). For these three datasets, we retained the first SNP of each RAD locus and we removed sites with minor allele frequency lower than 0.01. Second we used Tassel 5.0 (Bradbury et al., 2007) to filter the corresponding three VCF files according to missing data, with two strategies for each dataset: in one case we retained only loci present in at least 75 % of the individuals and individuals with data for 75 % of the loci (i.e. less than 25 % of missing data; hereafter strategy 75-75), in the other case we retained loci present in at least 95 % of the individuals, and individuals with data for 75 % of the loci (strategy 95-75). These filtering led to the removal of the less informative samples. The 75-75 strategy allowed to retain more loci with less individuals compared to the 95-75 strategy (Table 2). The final sampling sizes per sampling site and mitochondrial lineage for the six datasets are indicated in Supplementary Table S1.

**Table 2:** number of SNPs and individuals for the different datasets. The locus threshold corresponds to the minimum proportion of available data among individuals to retain a locus. The individual threshold corresponds to the minimum proportion of available data among loci to retain an individual. The mean depth indicates the mean depth per individual averaged over all individuals in the dataset. The last column indicates the number of retained individuals after correction for the presence of multiple MLL (see text for details). All corresponds to all samples (Polynesia and Oman).

| Dataset | Populations | Locus threshold | Individual threshold | Mean depth | SNPs | Individuals before MLL correction | Individuals after MLL correction |
| --- | --- | --- | --- | --- | --- | --- | --- |
| All_95_75 | All | 0.95 | 0.75 | 389 | 320 | 132 | 100 |
| All_75_75 | All | 0.75 | 0.75 | 66.9 | 194370 | 98 | 78 |
| Oman_95_75 | Oman | 0.95 | 0.75 | 155 | 1711 | 99 | 82 |
| Oman_75_75 | Oman | 0.75 | 0.75 | 72.7 | 134307 | 77 | 62 |
| Polynesia_95_75 | Polynesia | 0.95 | 0.75 | 433.3 | 558 | 31 | 29 |
| Polynesia_75_75 | Polynesia | 0.75 | 0.75 | 204.2 | 3285 | 25 | 18 |

### Population genetic analyses

We used the R package of the GENEPOP software (Rousset, 2008) to compute gene diversity within individuals (*1-Qintra;* corresponding to observed heterozygosity) and among individuals within samples (*1-Qinter;* corresponding to expected heterozygosity), and *F_IS_* (Weir & Cockerham, 1984). VCFTOOLS 0.1.15 (Danecek et al., 2011) was used to calculate an estimate of inbreeding coefficient, *F*, which compares the observed number of homozygous sites to its expectation under panmixia.

We tested the presence of repeated MLGs and multilocus lineages. Multilocus lineages (MLLs) correspond to genotypes separated by a varying number of mutations and potentially reflecting apparent divergence among MLGs either because of sequencing errors, ascertainment bias of individual SNPs to allelic states or because of somatic mutation since the last meiosis. We used the R package poppr to analyse MLGs and MLLs (Kamvar et al., 2015). The choice of thresholds to delineate MLLs was made according to two criteria: first we used MLGs obtained with microsatellite loci (data not shown) for a subset of individuals to define an MLL threshold. We also used the distribution of genetic distances among individuals to look for lowly differentiated individuals that could correspond to the same MLL. The genetic distances among individuals were measured by the percentage of nucleotidic divergence, and computed with poppr. According to their respective levels of diversity, the retained MLL threshold was different for the different datasets (see results). Networks based on the aforementioned distances were built with the NeighborNet option of SPLITSTREE 4.14.8 (Huson & Bryant, 2006). One representative of each MLL was kept for clustering and *F_ST_* analyses. We analysed the genetic disequilibrium among loci by computing the modified index of association 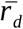 defined by (Agapow & Burt, 2001) with the poppr R package. To keep reasonable computing time, we first randomly subsampled the All_75_75 and Oman_75_75 to 25 000 SNPs. Then we computed 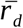 on datasets comprising randomly subsampled 200 SNPs (this number allowed enough different resampling with the smallest dataset), and with 10 000 repetitions of this subsampling. With this approach we can analyse the linkage disequilibrium in datasets with a high number of SNPs, where pairwise methods would be difficult to analyse. To take into account the impact of Wahlund effect on this analysis, we performed it at two levels: first at the level of the whole corresponding dataset, and second at the level of mitochondrial lineages for the All and Oman datasets, or of sampling sites for the Polynesia datasets (i.e. the “strata” levels used in poppr).

Genetic differentiation among populations was measured with the *F_ST_* estimator of Weir & Cockerham (1984) as computed with VCFTOOLS. The differentiation among individuals was visualised thanks to a Principal Component Analysis (PCA) with the R package adegenet (Jombart, 2008). Missing data were replaced by the mean allele frequency. As a complementary analysis to PCA, in order to identify the main genetic groups in the dataset, we analysed the partition in K independent units with the snmf function of the R package LEA (Frichot & François, 2015). This approach performs a least squares estimates of ancestry proportions (Frichot et al., 2014). We tested K values from 1 to 10, with ten replicates for each K value.

### Simulations

To help the interpretation of our results on individual inbreeding coefficient *F*, on 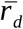 and on *F_ST_*, we performed simulations to analyse the behavior of these estimates, with a focus on the impact of partial clonality. We used SLiM 3 to build genetically explicit individual-based simulations (Haller & Messer, 2019). We simulated two populations, each with 100 individuals, and connected through reciprocal gene flow at a rate of 0.01 per generation. The genetic data were modeled with 2 000 loci of 100 bp each, mutating at a rate of 10^-4^ mutation per site per generation. This high mutation rate is a way to model enough genetic diversity but with a moderate number of individuals and memory usage. After an initial panmixia initialization phase of 5 000 generations, we performed 50 000 generations with one of the following reproductive mode: panmixia, clonality at a rate of 0.1, 0.5 or 0.9, selfing at a rate of 0.1, and a combination of 0.1 selfing rate and clonality rates of 0.1 or 0.5. At the end of the simulations, 30 simulated individuals were sampled, and 30 replicates were performed for each simulation configuration. The output VCF files were analysed with VCFTOOLS to compute the estimate of individual inbreeding coefficient *F* and *F_ST_*. Whenever possible, we computed *F_ST_* by comparing sampling sites or regions (Polynesia vs Oman), and by comparing individuals grouped according to the corresponding mitochondrial lineages. For each simulation we computed the mean, minimum and maximum values of *F* and *F_ST_* over individuals and loci respectively. We computed 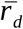 separately on each of the two population of the simulations. For computing reasons, the mean and standard deviation of 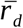 were computed with 50 resamplings of 1 000 SNPs.

## Results

### Assignment to species hypotheses according to mitochondrial sequences and microsatellite genotypes

Out of the 140 individuals retained in the final RAD-Seq dataset, we did not get any usable mitochondrial sequence for 18 individuals (three from Polynesia and 15 from Oman). We indicate in Supplementary Table S1 the correspondence between the nomenclature of mitochondrial lineages used in main text, the ORF haplotype number and the PSH and SSH as defined in Gélin *et al.*, 2017. The sequences obtained for the other individuals allowed a clear assignment to previously defined sequence groups and corresponding primary species hypotheses. In Polynesia, two individuals sampled as outgroups on the basis of morphology were highly divergent from other ones with RAD-Seq, and corresponded to mitochondrial lineages type 1a (PSH09 and SSH09c) and type 2 (PSH01). The high divergence of these individuals from other samples blurred the analysis of the differentiation among the other lineages, especially on multivariate analyses (data not shown). Regarding this signal and the small sample size for these outgroups, we did not retain them in the following analyses. Apart from these two individuals, Polynesia included only samples from mitochondrial lineage 5a (PSH5), and Oman included samples from mitochondrial lineage 7a (PSH12) and SSH13a, mitochondrial lineages 3g and 3e (both in SSH13a), and only one individual from mitochondrial lineage 5a (Supplementary Table S2). These assignments were confirmed with microsatellite loci (data not shown).

On the basis of microsatellites, we did not detect any repeated multilocus genotype (MLG) in Polynesia: each individual did correspond to a unique 13 loci genotype. However, among individuals from mitochondrial lineage 7a, over 64 individuals for which we got a 13 loci genotype, 53 distinct MLGs were retrieved implying one MLG repeated five times, another one four times in O2, and three MLGs repeated twice (one in O2 and two in O5). Over 20 genotyped individuals assigned to SSH13a, one MLG was repeated two times in O5.

### RAD sequencing data

The initial number of sequences obtained per individual was very uneven among samples: it varied from 5 735 to 30 394 029 reads (Supplementary Table S2). The mean number of read per individual was higher for samples from Oman (mean 5 647 233) compared to Polynesia (mean 2 309 860).

The percentage of reads aligned to the *Pocillopora* genome was more regular, with a mean of 85.1 %, but it was still higher in Oman (mean 78.3%) than in Polynesia (70.6%; Table S3). This heterogeneity among samples and sampling regions motivated our different assembly strategies. Table 2 presents the characteristics of the six final datasets: a total of 140 samples were distributed among the different datasets. The highest numbers of SNPs, with one SNP per RAD locus, were obtained for the 75 % filtering on loci missing data in the All (194 370 SNPs) and Oman datasets (134 307 SNPs). The separate assembly of Oman and Polynesia allowed the recovery of more SNPs than the All assembly with the 95_75 strategy, and a reverse result was obtained with the 75_75 strategy. The All_95_75 dataset had the highest number of individuals (132) and the lowest number of SNPs (320; Table 2).

### Genetic differences among individuals and repeated MLLs

The networks based on pairwise distances among individuals are presented in Figure 1 for the 95_75 datasets which had the highest number of individuals. The corresponding histograms of the distribution of genetic distances are presented in Supplementary Material (Figure S1). The All_95_75 network shows a clear distinction between Oman and Polynesia individuals, which is correlated to a difference in mitochondrial lineage: type 5a versus all other lineages, with one exception of type 5a sampled in Oman and grouping with other individuals from Oman. This distinction is also visible in the bimodal distribution of genetic distances (Figure S1). In the separate analysis of Oman and Polynesia, no clear sub-grouping appeared according to sampling location or mitochondrial lineages (Figure 1). Similar results were obtained for the networks based on the 75_75 datasets, except in Polynesia where we observed small clusters of individuals from the same site, albeit without clear differentiation from other groups (Supplementary Figure S2).

**Figure 1:**
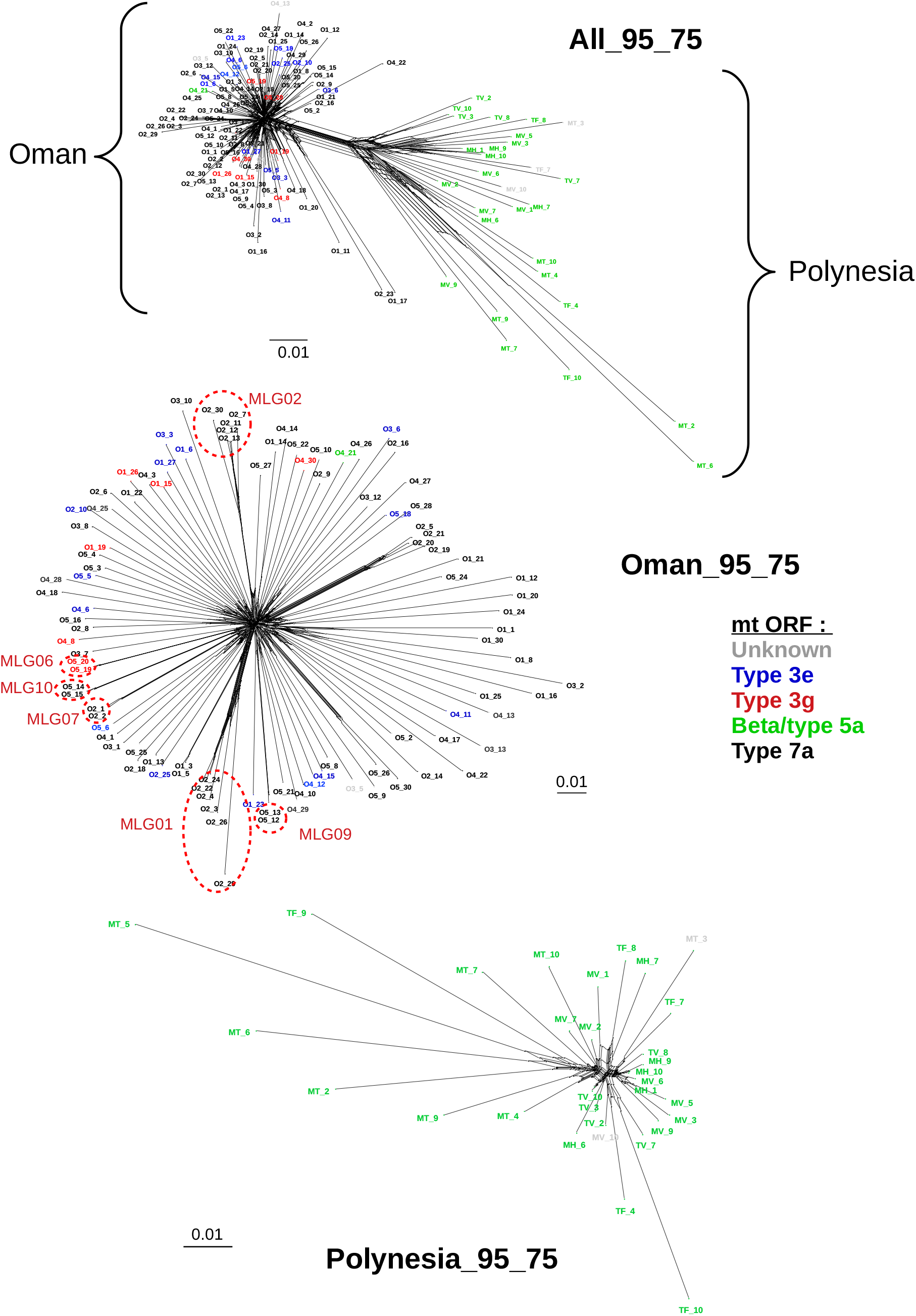
network based on the percentage of difference among individuals for the All, Oman and Polynesia 95_75 datasets. The colors indicate the corresponding species hypothesis according to the sequence of mitochondrial ORF. For clarity reasons we did not indicate all individual names for the All and Oman plots. The red ellipses indicate groupings of individuals corresponding to the same MLG according to microsatellite data.

The histograms of pairwise differences among individuals in Oman, and to a lesser extent in Polynesia, showed a peak of low value distances, potentially reflecting repeated MLLs. We used the distance observed with RAD-Seq among repeated microsatellite MLGs to define MLLs in the Oman dataset: we analysed the distance among all individuals corresponding to identical microsatellite MLGs. We estimated the distance (percentage of differences) with SNPs for individuals with identical microsatellite MLG. We retained the second highest of these distance as a threshold to define MLLs with SNPs, because the highest distance among individuals with identical microsatellite MLGs seemed too high to be used to define MLLs with SNPs. We could not use a single threshold for all datasets because the levels of divergence were very different between the datasets (95_75 vs 75_75, and Polynesia vs Oman), and we had no repeated microsatellite MLG to be used as a reference in some datasets (see above). Therefore, in cases where no threshold could be defined on the basis of microsatellites, we used a threshold allowing the removal of the closest individuals, as indicated by preliminary tests and by the observation of the distribution of pairwise distances among individuals. The number of individuals for each corrected dataset is given in Table 2, and the corresponding thresholds are indicated below the distribution of individual distances (Figure S1).

### Genetic diversity and differentiation

The parameters of genetic diversity for the different datasets are presented in Table 3. Separate estimates of genetic diversity per lineage or site are presented in Supplementary Material (Table S4). The estimates of genetic diversity were higher in the 75_75 datasets compared to the 95_75 ones. The mean genetic diversity within individuals (‘1-Qintra’) varied between 0.04 in Polynesia_95_75 and 0.20 in Oman_75_75. The mean genetic diversity among individuals (‘1-Qinter’) varied between 0.04 in Polynesia_95_75 and 0.24 in Oman_75_75. The *F_IS_* were mainly positive in the 75_75 datasets and negative to positive in the 95_75 ones, but with important variations among lineages or sites, especially in Polynesia (Table S4). The estimates of individual inbreeding coefficient *F* gave highly variable and extreme values (Table 3): the lowest minimum values were observed in the 95_75 datasets, and in Polynesia (down to −2.733 in Polynesia_95_75), which indicated the presence of individuals with an excess of hetezygous loci. The maximum *F* value was observed in the All_75_75 dataset (0.732). The distributions of the *F* estimates illustrate this wide dispersion, and the shift to more positive values from 95_75 to 75_75 datasets (Figure S3). In the All_95_75 dataset, the *F* values in Oman (from −0.278 to 0,546) were higher than in Polynesia (from −1,306 to −0.303). We analysed whether the individuals involved in potential MLLs (i.e. individuals involved in the closest pairwise relationships in the different datasets) corresponded to particularly high or low *F* values (Figure S3). In the All datasets, the closest individuals showed among the highest *F* values for the 95_75 dataset, whereas for the 75_75 dataset they also included the lowest *F* values. For the Oman datasets, these individuals showed a wide range of *F* negative and positive values. For the Polynesia datasets, these individuals were among the highest *F* values. The results of the analysis of linkage disequilibrium with the 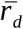 index are presented in supplementary material (Table S5 and Figure S4). For the analyses at the level of the whole datasets, the highest 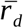 values were obtained in the All_75_75 dataset, followed by the Polynesia_95_75 and All_95_75 datasets. The values obtained in Polynesia were higher than those in Oman for the 95_75 and the 75_75 datasets. When the analysis was performed at the level of mitochondrial lineages or sites, the highest values were observed for the MH site in Polynesia with mean values of 0.278 for the 95_75 dataset, and 0.235 for the 75_75 dataset (Table S4, Figure S4).

**Table 3:**
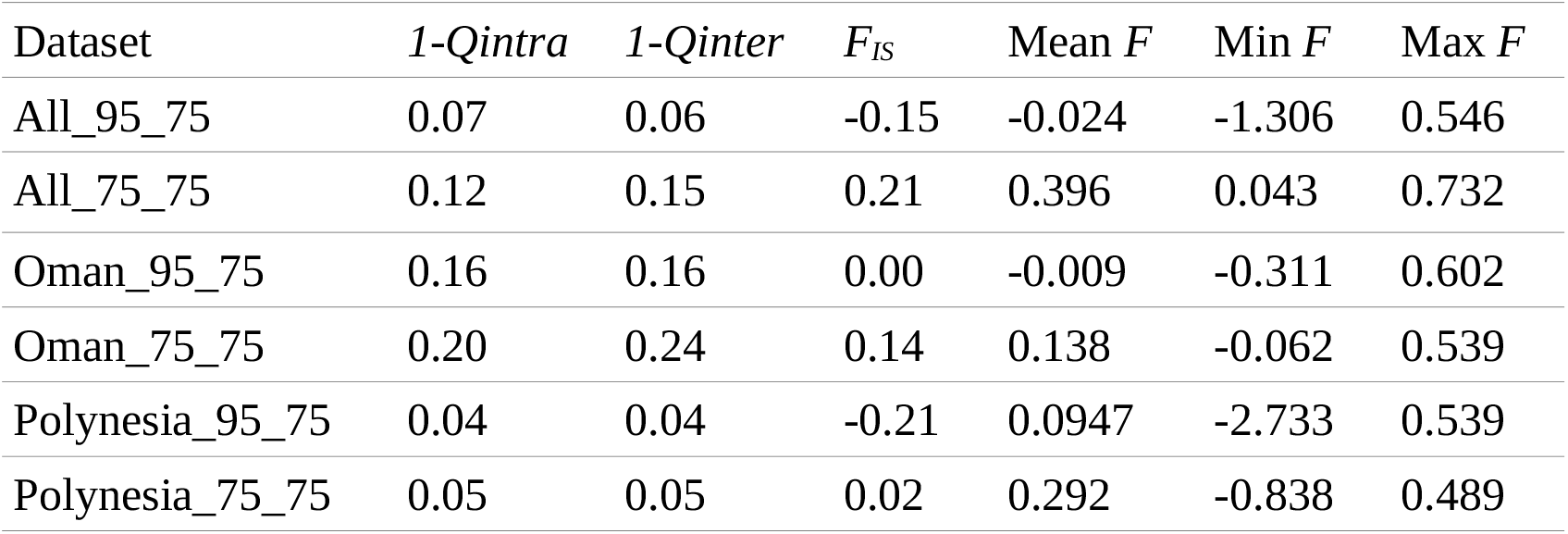
estimates of gene diversity within (*1-Qintra*) and among individuals (*1-Qinter*), and of *F_IS_* averaged over samples. For the All datasets, the average was done over Oman and Polynesia. For the Oman and Polynesia datasets, average was done over sampling locations. The last three columns provide indicators of the distribution of inbreeding coefficient (mean, mininum and maximum) computed over all individuals for each dataset.

The mean *F_ST_* estimates among loci were generally lower for the datasets corrected for MLLs compared to the non-corrected datasets (Table 4), except for the comparison among mitochondrial lineages with All_95_75, and the comparison among populations with All_75_75. The mean *F_ST_* between Polynesia and Oman (by grouping samples from each region) was 0.105 and 0.352 for the All_95_75 and the All_75_75 datasets respectively. The distributions of *F_ST_* among loci for these Oman / Polynesia comparison are presented in Figure S5. The distribution of *F_ST_* estimates was very different between the two datasets, despite a peak around 0 in both cases: for All_75_75, there was an important proportion of loci with *F_ST_* above 0.2, and a peak at *F_ST_* = 1, whereas the distribution was mainly restricted to values below 0.2 for All_95_75. For Oman, the 75_75 *F_ST_* estimates were slightly higher than the 95_75 ones, but in all cases (75_75 or 95_75, population or mitochondrial lineages comparisons) these estimates indicated very low levels of differentiation in Oman, with values ranging from −0.006 (Oman_95_75 for the comparison of mitochondrial lineages, with MLL correction) to 0.016 (Oman_75_75 for the comparison of populations, no correction; Table 4). The Polynesia *F_ST_* estimates also indicated very low levels of differentiation among sampling sites. Regarding multivariate analyses, the PCA of the All_95_75 and All_75_75 datasets separated the samples from Polynesia and Oman, but with individuals from Polynesia (type 5a lineage) more spread apart than those from Oman (Figure S6). The separate PCAs on the Oman and Polynesia datasets did not reveal any clear structure according to sampling site nor mitochondrial lineage, whatever the filtering strategy.

**Table 4:** mean *F_ST_* estimates among loci for the datasets without and with correction for repeated MLLs. The “pop” comparison estimates *F_ST_* by comparing sampling sites; for the All dataset, it corresponds to the Polynesia / Oman differentiation. For the “ORF” comparison, this estimate compares samples of individuals grouped according to their ORF haplotypes (individuals without ORF sequence were not taken into account).

| Dataset | $F_{ST}$ all individuals | $F_{ST}$ corrected for MLLs |
| --- | --- | --- |
| All_95_75 pop | 0.105 | 0.088 |
| All_95_75 ORF | 0.178 | 0.186 |
| All_75_75 pop | 0.352 | 0.355 |
| All_75_75 ORF | 0.361 | 0.312 |
| Oman_95_75 pop | 0.011 | 0.001 |
| Oman_95_75 ORF | 0.003 | -0.003 |
| Oman_75_75 pop | 0.016 | 0.002 |
| Oman_75_75 ORF | 0.005 | -0.002 |
| Polynesia_95_75 pop | 0.000 | -0.007 |
| Polynesia_75_75 pop | 0.015 | -0.024 |

The analysis of genetic structure with the snmf method did not give a clear signal for an informative K value on the basis of cross-entropy plots, apart for the All_95_75 dataset where a first minimum value was observed at K = 2 (and a second at K = 5), and for Polynesia_95_75 where a minimum was observed at K = 2 (data not shown). We also analysed the results corresponding to the number of mitochondrial lineages (for All and Oman) or the number of sampling sites (for Polynesia) as the retained K value. The corresponding barplots of coancestry coefficients for the 95_75 datasets are given in Figure 2. For All_95_75, the K = 2 solution clearly separated Polynesia (mitochondrial lineage type 5a) and Oman samples (other mitochondrial lineages, with one 5a exception). At K = 4, two additional sub-clusters were observed, one in Oman and one in Polynesia. In Polynesia this new structuring did not clearly separate the sampling sites. In Oman the sub-cluster grouped five individuals with the mitochondrial lineage 7a from the O2 sampling site: these individuals corresponded to the same MLG identified with microsatellites and confirmed by the proximity of individuals on the individual network (group MLG02 in the Oman_95_75 network, Figure 1). For the Oman_95_75 dataset, the K = 4 solution led to a major and three minor clusters. These clusters did not separate individuals neither by mitochondrial lineage nor by sampling site. This clustering was nevertheless partly linked with potential MLLs: the purple cluster of Oman_95_75 grouped individuals of the MLG02 identified with microsatellites (see Figures 1 and 2). For Polynesia_95_75 the K = 2 solution separated three individuals which did not correspond to a potential MLL. The K = 5 solution for Polynesia_95_75 led to a major and four minor clusters: this clustering did not separate individuals neither by sampling site type nor by potential MLL. The snmf analysis of genetic structure with the 75_75 datasets gave similar results, with a separation of Oman and Polynesia at K = 2 for All_75_75, and a further distinction of a few individuals in two additional clusters in Oman at K = 4 (results not shown).

**Figure 2:**
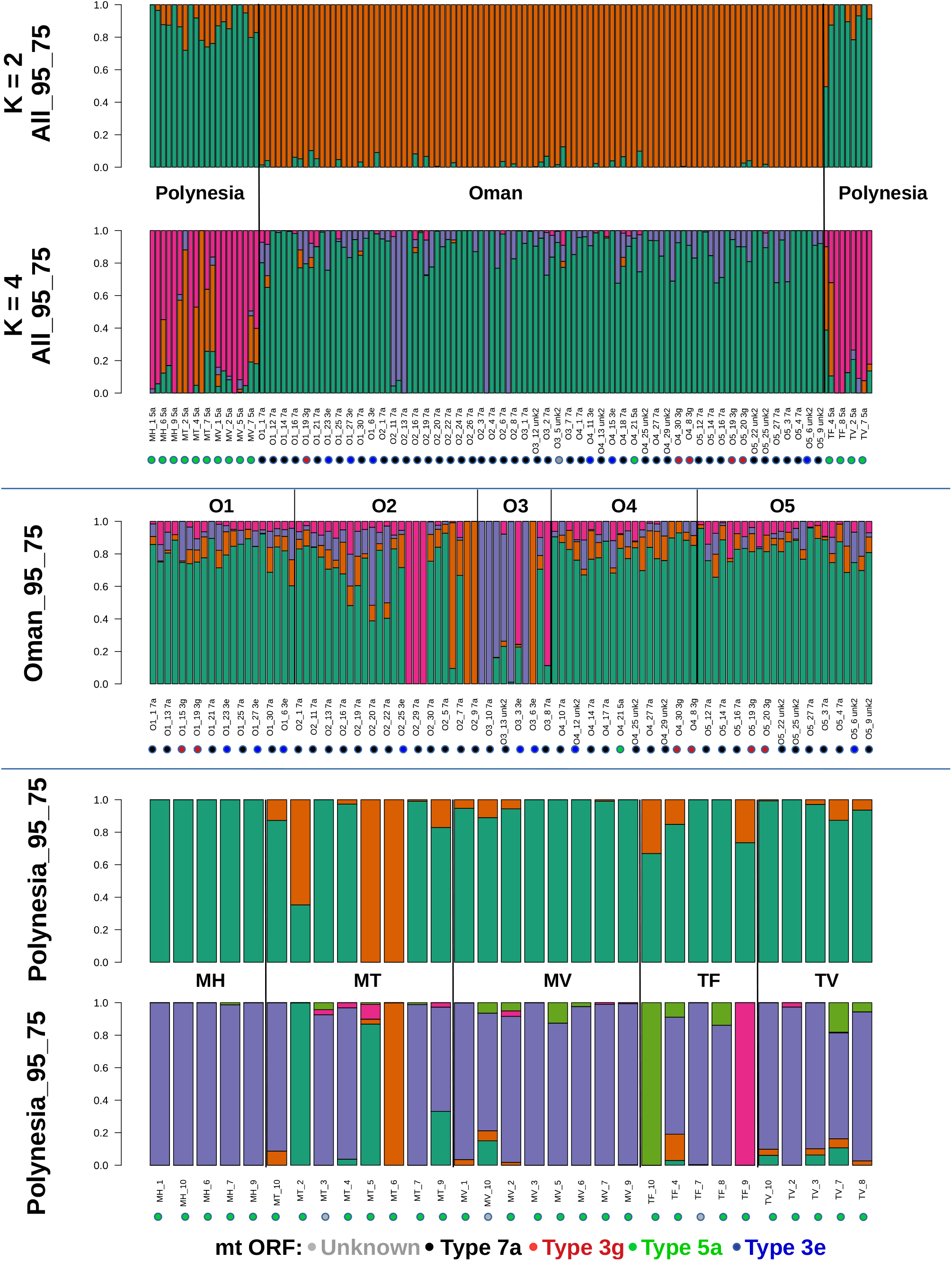
plots of coancestry coefficients inferred with the LEA R package for the 95_75 datasets. For the All and Polynesia datasets, the cross-entropy gave a first minimum value at K = 2. Apart from these cases we present here the results obtained by considering the number of ORF mitochondrial lineages in the dataset (K = 4 for All and Oman), and the number of population (K = 5 in Polynesia). Color dots under individual names indicate the ORF mitochondrial lineage. For clarity reasons, only legends of one individual over two are written under the All and Oman plots.

### Simulations

The distribution of the individual inbreeding coefficients *F*, of the indices of association 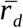, and of the estimates of *F_ST_* for the different simulation configurations are presented in Supplementary Material. The average and *F* values were higher for the simulations including selfing. The maximum *F* values were highly variable and tended to be higher for the configurations with the highest clonality rates (from 0.5), but were much higher for the simulations including selfing. Regarding the minimum *F* value, a decrease in the distribution was observed for the highest levels of clonality compared to other configurations. When comparing these results with observed data, one should note that for the All dataset, the *F* estimates were based on the pool of Oman and Polynesia samples, whereas for the simulations we analysed the two simulated populations separately. The average *F* obtained in the All_75_75 and Polynesia_75_75 was higher than all values obtained with simulations. Regarding the maximum *F* values, the highest values observed in All_75_75 and Oman_95_75 appeared only compatible with simulations integrating selfing. For the minimum *F* values, no configuration allowed to recover such highly negative *F* as those observed in Polynesia (both datasets), nor in All_95_75.

We analysed the linkage disequilibrium within each of the two simulated populations with the 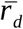 index. An increase in 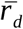 was observed with increasing clonality rate, mainly for the highest clonality rate tested here (0.9). A much higher increase both in mean and standard deviation was observed for a combination of selfing and a 0.9 clonality rate. We compared these values to the highest observed values obtained with the 75_75 datasets. Apart from one value in Oman, these mean observed values were similar or higher to those obtained with simulations at clonal rates of 0.9. The standard deviation for the observed estimates were usually very high, and in most cases these values were only approached by simulations with selfing and a 0.9 clonal rate.

Regarding the average *F_ST_*, without any variation neither in census size nor migration rate, the resulting values were mostly similar among simulation configurations. A slight decrease and higher variance was nevertheless observed for the highest clonality rate (0.9). The observed *F_ST_* for the Oman and Polynesia datasets were lower than those obtained in all simulations, while the *F_ST_* for the All datasets were higher than those obtained in almost all simulations, except a for a few simulations performed with the highest clonality rates.

## Discussion

### Impact of filtering strategies on RAD-Seq data

We obtained very uneven levels of missing data in our results. High levels of missing data, if not accounted for, can lead to incorrect conclusions regarding genetic structure for example (Larson et al., 2021). Missing data in RAD-Seq can have several origins including mutations in enzyme-cutting sites, technical problems linked to library preparation, uneven amplification or sequencing, or errors in the *in silico* identification of homologous sites (Eaton et al., 2017; O’Leary et al., 2018). Here, the lowest read numbers were obtained for outgroup samples, corresponding here to species hypotheses PSH01 and SSH09c (as opposed to PSH05, PSH12 and SSH13a which were the most frequent in our datasets): for example seven of these samples had only around 20 000 reads or less and were not retained (data not shown). Such very low levels of reads rather points to difficulties in library preparation. The laboratory protocol is not into question, as we used it on the octocoral *Corallium rubrum*, and we obtained much more regular results (Pratlong et al., 2018). We also did not observe any relationship between the number of raw reads and the number of reads mapping to a *Symbiodinium* genome (data not shown): therefore a contamination from dinoflagellate genomes cannot explain these results. Despite standard verifications, problems with DNA quantity and quality may have impacted the number of reads, such as for example partially degraded DNA or the presence of inhibitors (O’Leary et al., 2018).

Facing these difficulties, we compared different strategies for filtering missing data, which can have important consequences on the obtained results. Some results were stable among the different datasets: there was a marked differentiation between samples from Oman and Polynesia, correlated with the species sampled in each region, and a lack of genetic structure inside each region. Conversely the estimates of genetic diversity and structure differed among datasets. The genetic distances among individuals were much higher with more loci in the All and Oman datasets.

If part of missing data are linked to mutations in the cutting site, filtering loci according to their rates of missing data is expected to reduce the frequency of loci with high mutation rates (Huang & Knowles, 2016), which would agree well with our observation of a lower diversity with more stringent filtering. Allele dropout, which corresponds to the non-observation of a SNP linked to a mutated restriction site, should be more frequent in the 75_75 datasets (with low filters on SNP missing data). Allele dropout is expected to lead to an overall underestimate of genetic diversity, increasing with the level of polymorphism (Arnold et al., 2013; Cariou et al., 2016). Contrarily to these expectations, we observed more diversity with less filtering on SNPs. For the retained variable loci, an overestimation of heterozygosity can be expected with allele dropout if they concern more ancestral allelic states and therefore lead to an increase in minimum allele frequencies (Gautier et al., 2013); accordingly we observed an increase in heterozygosity for the 75_75 compared to the 95_75 datasets. We also observed an increase of *x* for the 75_75 compared to the 95_75 datasets, but the reason of this effect remains to be studied: one possible explanation could be a difference in the sampling of clonal lineages depending on datasets (see below). Finally, allele dropout can overestimate *F_ST_* (Gautier et al., 2013) which seems coherent with our results (with higher *F_ST_* for the 75_75 compared to the 97_75 All datasets). Regarding linkage disequilibrium, the increase in 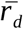 observed for All75_75 compared to All_95_75 can result from a combination of higher Wahlund effect (with the inclusion of more differentiated loci), and of a higher number of physically linked loci.

Importantly, the potential impact of missing data and allele dropout on our results is potentially blurred by the fact that these filtering led to different numbers of retained individuals. Retaining less variable loci in the 95_75 datasets may have favored the retention of less divergent individuals which can impact estimates of relatedness or clonality. Another question is whether the variance of polymorphism among loci is high enough to explain the observed differences through the aforementioned effects. Along with allele dropout, other effects can take place to create differences of diversity among datasets. For example paralogs and copy number variants (CNV) were probably differently represented among datasets: a perspective of this study would be to use RAD-Seq data to study CNV loci through the use of proportion of heterozygous and read ratio deviation of alleles (Dorant et al., 2020; McKinney et al., 2017).

### Signals of clonality with RAD-Seq data

Populations of *Pocillopora* corals, notably in PSH05 (*P. acuta*) or PSH04 (*P. damicornis*), can show various levels of clonal reproduction (Adjeroud et al., 2014; Pinzón et al., 2012; Torda et al., 2013), with sometimes different ramets of the same genet separated by several kilometres (Gélin et al., 2017, 2018). Clonal reproduction in these species can happen through fragmentation of individuals, polyp bail-out or asexual production of larvae (Gélin et al., 2017; Highsmith, 1982; Oury et al., 2019). Clonal reproduction can lead to heterozygote excess compared to panmixia (Balloux et al., 2003; Reichel et al., 2016), and to shift the distribution of *F_IS_* among loci towards negative values for the highest rates of clonality (Stoeckel & Masson, 2014). This is well visible in strictly asexual organisms such as the Euglenozoa *Trypanosoma brucei gambiense* (Weir et al., 2016). Despite the difficulty to delineate clonal lineages, RAD-Seq offers new avenue in the study of the reproduction of partially clonal organisms, such as in the Kelp *Laminaria rodriguezii* (Reynes et al., 2021). Several points in our results show the effect of clonality in the studied populations. First, samples corresponding to MLGs detected with microsatellites were indeed grouped with reduced distance in networks based on RAD-Seq. Second, the distribution of pairwise differences showed a peak of low divergence which can be an indication of repeated MLLs. Third, the distribution of estimates of the inbreeding coefficient *F* showed some individuals with very negative values, corresponding to very heterozygous individuals, especially in Polynesia. Our results point to the first observation of clonality in SSH13/PSH12, i.e. *P. verrucosa* / mitochondrial lineage 7a (no repeated microsatellite MLGs found over thousands of PSH13 individuals in Oury *et al.*, 2021). This could be explained by the fact that the *P. verrucosa* individuals were sampled in shallow water, with a *corallum* macromorphology similar to *P. acuta* morphology (i.e. thin branches highly breakable). So this pattern could rather be explained by the environment where they have been sampled, and where they might be submitted to waves and swell, favoring fragmentation and thus clonal propagation. We also observed individuals with very low *F* values, indicating high rates of heterozygous loci. This was observed in our simulations only with the highest clonal rates which are not compatible with the low frequency of repeated MLLs observed here. Other effects could explain these observations, such as the presence of brooded larvae, or the intra-colonial genetic diversity linked to chimerism or mosaicism (Schweinsberg et al., 2015). Hybrids between divergent lineages could also create such high heterozygosity, but could not be detected here if the corresponding lineages were not analysed. In all cases estimating the rate of clonality in these populations would require a dedicated sampling.

From a methodological point of view, our simulations provide new avenues in the study of clonality with RAD-Seq data. Our simulations showed a discernible effect of clonality on the distribution of *F* and 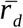 for the highest clonality level tested here (0.9). This is in line with previous studies demonstrating an effect of clonality only for extreme rates of clonality (Balloux et al., 2003). Though, the scope of our simulations is limited on several points. First, we did not explore the impact of sampling scheme on the estimates of genetic diversity and genetic structure. Second, *Pocillopora* corals show overlapping generations, and a given clone may persist over several generations, which was not possible in our simulation framework. Third, selective effects can lead to the expansion of one clone (see Gélin et al., 2017) and references therein), and modify the distribution of clones. These limits evidently hinder estimates of clonal rates based on these simulations. This would require simulations taking into account these limits, for example coupled with Approximate Bayesian Computation (Csilléry et al., 2010). We should also take into account the robustness of these results to varying sample sizes: for example 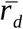 is based on the pairwise distances between individuals, therefore it may be sensitive to low sample sizes (as was the case here for some empirical datasets).

Regarding genetic structure, our simulations showed a slight decrease of *F_ST_* only for a clonal rate of 0.9: this agrees well with theoretical expectations (Balloux et al., 2003). The correction of datasets for repetitions of clonal lineages changed the estimated *F_ST_* values but this did not change our main conclusions: an important differentiation for the All datasets, and low to no genetic structure in Oman and Polynesia. In some cases such correction for clonal diversity can lead to very different conclusions, from genetic differentiation without correction to near panmixia in *Pocillopora* in French Polynesia (Adjeroud et al., 2014). Therefore, one should not rely on a single analysis strategy (e.g. all individuals or single genotypes for example), but consider the different results thus obtained (De Meeûs et al., 2006).

### Genomic analysis of species hypotheses

When considering the All datasets, we observed a marked differentiation between Polynesia and Oman which was superimposed on a differentiation between PSH05 and other species hypotheses (with one exception in Oman). This was well visible on the network, PCA and snmf analyses, both with the 97_75 and the 75_75 filtering strategies. The mitochondrial lineage 5a corresponds to the PSH05, and is phylogenetically well separated from lineages 3e −3g and 7a which correspond to SSH13a and PSH12 (Gélin et al., 2017). The distinction of PSH05 from PSH 12 and SSH13a was previously confirmed with microsatellite loci (Gélin et al., 2017). Our results confirm the distinction of PSH05, but our sampling scheme did not allow to test the delimitation and possibility of hybridization of this species with other species in sympatry. There was only one individual bearing mitochondrial type 5a in Oman for which we were able to get RAD-Seq data: this individual did not separate from other individuals in Oman with different mitochondrial types. This could indicate a possible introgression of type mitochondrial 5a in the 3e-3g-7a gene pool, but this should be tested by considering additional 5a individuals in Oman. We obtained six additional individuals from this lineage, hence indicating that the single individual reported here is not likely a contamination, but the RAD sequencing of these samples was not good enough to retain them. This discrepancy between RAD-Seq and microsatellites, if confirmed, could point to a genomic heterogeneity in introgression following secondary contact, or to an effect of considering a single PSH05 individual with RAD-Seq.

Conversely, we did not observe a differentiation between individual from Oman assigned to the species hypotheses PSH12 and SSH13a according to their mitochondrial haplotype. All methods of species delimitation based on mitochondrial ORF used in Gélin et al. (2017) indeed separated these two species hypotheses, whereas this was not the case with microsatellites in this previous article and in the current one. With a study in sympatry at a genomic scale, we also reject this species delimitation based on mitochondrial sequences. The most parsimonious hypothesis here would be that these lineages correspond to mitochondrial polymorphism present in a given species, here *P. verrucosa*, even if mitochondrial DNA in anthozoans has been shown to evolve slower than in other metazoans (Calderón et al., 2006; Hellberg, 2006; van Oppen et al., 1999). Accordingly, one can note that 12 over 16 species delimitation methods based on mitochondrial ORF did not conclude to separate PSHs for the lineages 3e and 3g (Gélin et al., 2017). Another possibility could be a genetic swamping (e.g. Bog et al., 2017) following a secondary contact between different lineages in Oman. For example, in the plant *Salix serpillifolia*, the discrepancy between the genetic structure observed with plastid microsatellites and the much lower structuring with nuclear ones, has been attributed to genetic swamping (Kosiński et al., 2019). Reticulate evolution has already been proposed as a major factor shaping the current diversity of scleractinian corals (van Oppen et al., 2001; Vollmer & Palumbi, 2002). A more precise analysis of genomic patterns of diversity and differentiation, would be useful to go further on these questions in *Pocillopora* spp. (e.g. (Nelson et al., 2020), and if possible with the use of high-quality genome (Manel et al., 2016). It would also be necessary to include additional PSHs to better estimate the range of genomic divergence among species in this genus.

In Polynesia, both datasets did not show evidence of genetic structure among sites, which were distributed in the two islands of Moorea and Tahiti. Though limited by a reduced number of individuals, these results are in line with previous studies on the genetic structure of *Pocillopora*: (Adjeroud et al., 2014; Magalon et al., 2004; Oury et al., 2021). One can note that in *Pocillopora* corals the patterns of genetic structure are evidently dependent on the species hypothesis and location considered (Oury et al., 2020).

## Conclusions and perspectives

Because of unequal success in the production of RAD sequences, we obtained a dataset with very different levels of missing data among samples, and potentially low number of loci shared among samples depending on the level of analysis. We therefore had to explore different filtering strategies to get informative datasets. The trade-off between the number of loci and individuals retained in the datasets led to different estimates of genetic diversity. Nevertheless the main patterns of genetic structure were conserved among the different datasets, which shows the interest of testing the impact of data filtering on the obtained results. Interestingly, we can conclude that the main signal of genetic structure was visible with only slightly more than 300 SNPs. Of course, depending on the objectives of the study, higher number of loci may be necessary. For example an analysis of outlier loci potentially under selection will require enough markers with good confidence in genotyping. Considering the aforementioned difficulties we did not develop such approaches here, but this would be interesting to develop studies on local adaptation and speciation processes in *Pocillopora* spp. This can be of interest to integrate evolutionary processes in management and conservation (e.g. Xuereb et al., 2020).

Our results on the divergence among Polynesia and Oman, and on the low genetic structure inside each region, were robust to the different strategies of analysis. We were also able to detect signals of clonal reproduction in the sampled populations. The study of clonality with reduced representation genomic data in populations of partially clonal species is not yet well developed. Through some comparisons with microsatellite data, simulations or among replicated samples, this will provide a powerful tool to the study of mixed reproduction systems.

## Acknowledgements

This work is a contribution to the Labex OT-Med (n° ANR-11-LABX-0061) funded by the French Government “Investissements d’Avenir” program of the French National Research Agency (ANR) through the A*MIDEX project (n° ANR-11-IDEX-0001-02). This project has been funded by the ADACNI program of the French National Research Agency (ANR) (project n°ANR-12-ADAP-0016; http://adacni.imbe.fr). The project leading to this publication has received funding from European FEDER Fund under project 1166-39417. We thank Nicolas Fernandez and Béatrice Loriod from the Marseille TGML platform for their invaluable help and advice with the preparation of the RAD libraries; the team of the MGX platform for the sequencing of the RAD libraries. The authors thank the UMR 8199 LIGAN-PM Genomics platform (Lille, France, especially Véronique Dhennin) which belongs to the ‘Federation de Recherche’ 3508 Labex EGID (European Genomics Institute for Diabetes; ANR-10-LABX-46) and was supported by the ANR Equipex 2010 session (ANR-10-EQPX-07-01; ‘LIGAN-PM’). The LIGAN-PM Genomics platform (Lille, France) is also supported by the FEDER and the Region Nord-Pas-de-Calais-Picardie. This study was set within the framework of the Laboratoire d’Excellence (LABEX) TULIP (ANR-10-LABX-41) We acknowledge the staff of the “Cluster de calcul intensif HPC” Platform of the OSU Institut Pythéas (Aix-Marseille Université, INSU-CNRS) for providing the computing facilities. We gratefully acknowledge Julien Lecubin and Christophe Yohia from the Informatic Service of Pythéas Institute (SIP) for their technical assistance. We acknolwedge Stéphanie Rialle, Marine Maurine Bonabaud from the MGX sequencing platform (CNRS, Montpellier, France) for the help with data production and quality controls. The authors acknowledge the financial support from the France Génomique National Infrastructure, funded as part of “Investissement d’Avenir” program managed by the Agence Nationale de la Recherche (contract ANR-10-INBS-09). We thank the molecular biology service of the IMBE.

## Data acessibility

Raw sequences are available in Genbank under BioProject ID PRJNA689941 and SRA accession number SRA PRJNA689941. The mitochondrial ORF sequences, microsatellite genotypes and SliM scripts are available in Zenodo: https://zenodo.org/record/4748346

The scripts used for SLiM simulations are also available at https://gitlab.osupytheas.fr/aurelle/slim-simulations.

