## Supplementary material for "Population genomics of *Pocillopora* corals: insights from RAD-sequencing"

**Table S1:** correspondence between the mitochondrial lineage nomenclature used in the manuscript ORF haplotypes, and primary and secondary species hypotheses (PSH and SSH respectively ; see Gélin *et al.*, 2017 for details). *Note that the correspondence between mitochondrial lineages and SSH indicated here for our samples is based on the complementary analysis of microsatellite genotypes: it may therefore be different for other samples.*

| Mitochondrial lineage | ORF | PSH | SSH |
|---|---|---|---|
| type 1a | 27 | 9 | 9c |
| type 2 | 1 | 1 | 1 |
| type 5a | 18 | 5 | 5a |
| type 7a | 34 | 12 | 12 |
| type 3e | 36 | 13 | 13a |
| type 3g | 43 | 13 | 13a |

**Table S2:** samples sizes per species hypothesis (on the basis of mitochondrial ORF) and sampling site for the different datasets. See main text for details on sampling sites and Gélin et al. (2017) for species hypotheses. For Polynesia, all mitochondrial sequences corresponded to type 5a. Note that some individuals were shared among datasets.

| **Dataset** | **Region** | **Site** | **type 5a** | **type 7a** | **type 3g** | **type 3e** | **unknown** |
|---|---|---|---|---|---|---|---|
| All_95_75 | Oman | O1 | | 16 | 3 | 3 | |
| All_95_75 | Oman | O2 | | 24 | | 2 | |
| All_95_75 | Oman | O3 | | 6 | | 2 | 2 |
| All_95_75 | Oman | O4 | 1 | 13 | 2 | 4 | 1 |
| All_95_75 | Oman | O5 | | 19 | 2 | 3 | |
| All_95_75 | Polynesia | MH | 5 | | | | |
| All_95_75 | Polynesia | MT | 6 | | | | 1 |
| All_95_75 | Polynesia | MV | 7 | | | | 1 |
| All_95_75 | Polynesia | TF | 3 | | | | 1 |
| All_95_75 | Polynesia | TV | 5 | | | | |
| **All_95_75** | | **Total** | **27** | **78** | **7** | **14** | **6** |
| All_75_75 | Oman | O1 | | 5 | 3 | 3 | |
| All_75_75 | Oman | O2 | | 19 | | 2 | |
| All_75_75 | Oman | O3 | | 5 | | | 2 |
| All_75_75 | Oman | O4 | 1 | 11 | 2 | 3 | |
| All_75_75 | Oman | O5 | | 19 | 2 | 3 | |
| All_75_75 | Polynesia | MH | 4 | | | | |
| All_75_75 | Polynesia | MV | 6 | | | | 1 |
| All_75_75 | Polynesia | TF | 1 | | | | 1 |
| All_75_75 | Polynesia | TV | 5 | | | | |
| **All_75_75** | | **Total** | **17** | **59** | **7** | **11** | **4** |
| Oman_95_75 | Oman | O1 | | 14 | 3 | 3 | |
| Oman_95_75 | Oman | O2 | | 23 | | 2 | |
| Oman_95_75 | Oman | O3 | | 7 | | 2 | 1 |
| Oman_95_75 | Oman | O4 | 1 | 13 | 2 | 4 | |
| Oman_95_75 | Oman | O5 | | 19 | 2 | 3 | |
| **Oman_95_75** | | **Total** | **1** | **76** | **7** | **14** | **1** |
| Oman_75_75 | Oman | O1 | | 5 | 3 | 3 | |
| Oman_75_75 | Oman | O2 | | 18 | | 2 | |
| Oman_75_75 | Oman | O3 | | 5 | | | 1 |
| Oman_75_75 | Oman | O4 | 1 | 10 | 2 | 3 | |
| Oman_75_75 | Oman | O5 | | 19 | 2 | 3 | |
| **Oman_75_75** | | **Total** | **1** | **57** | **7** | **11** | **1** |
| Polynesia_95_75 | Polynesia | MH | 5 | | | | |
| Polynesia_95_75 | Polynesia | MT | 7 | | | | 1 |
| Polynesia_95_75 | Polynesia | MV | 7 | | | | 1 |
| Polynesia_95_75 | Polynesia | TF | 4 | | | | 1 |

| | | | | |
|---|---|---|---|---|
| Polynesia_95_75 | Polynesia | TV | 5 | |
| **Polynesia_95_75** | | **Total** | **28** | **3** |
| **Polynesia_75_75** | Polynesia | MH | 5 | |
| **Polynesia_75_75** | Polynesia | MT | 6 | 1 |
| **Polynesia_75_75** | Polynesia | MV | 7 | 1 |
| **Polynesia_75_75** | Polynesia | TV | 5 | |
| **Polynesia_75_75** | | **Total** | **23** | **2** |

**Table S3 :** statistics on read numbers and alignment after initial filtering on sequence quality : mean, minimum, maximum and standard deviation (s.d.) of the initial numbers of reads, and of the percentage of reads mapped on genome par individual.

| Samples | Number of individuals | Mean reads | Min reads | Max reads | s.d. | Mean mapped | Min mapped | Max mapped | s.d. |
|---|---|---|---|---|---|---|---|---|---|
| All | 211 | 4540048 | 5735 | 30394029 | 5169991 | 85.1 | 70.6 | 86.7 | 1.9 |
| Oman | 141 | 5647233 | 5735 | 30394029 | 5669214 | 85.6 | 78.3 | 86.7 | 1.1 |
| Polynesia | 70 | 2309860 | 20495 | 11800706 | 2930211 | 84.1 | 70.6 | 86.2 | 2.7 |

**Table S4 :** estimates of gene diversity within (*1-Qintra*) and among individuals (*1-Qinter*), and of $F_{IS}$ over all loci. For each dataset the estimates are given per population (here sampling site) and per ORF haplotype if at least two individuals with the corresponding ORF haplotype have been analysed. The results are given for datasets not corrected for repeated MLLs. For Polynesia, all individuals for which we got a mitochondrial ORF sequence corresponded to beta/type5a.

| **Dataset** | **Population / ORF** | ***1-Qintra*** | ***1-Qinter*** | $F_{IS}$ |
|---|---|---|---|---|
| **All 95 75** | | | | |
| | **Population** | | | |
| | Polynesia | 0.09 | 0.09 | -0.04 |
| | Oman | 0.05 | 0.04 | -0.27 |
| | **ORF** | | | |
| | type5a | 0.09 | 0.09 | -0.01 |
| | 7a | 0.05 | 0.04 | -0.27 |
| | 3g | 0.04 | 0.03 | -0.42 |
| | 3e | 0.05 | 0.04 | -0.28 |
| **All 75 75** | | | | |
| | **Population** | | | |
| | Polynesia | 0.13 | 0.17 | 0.27 |
| | Oman | 0.11 | 0.13 | 0.15 |
| | **ORF** | | | |
| | type5a | 0.13 | 0.19 | 0.33 |
| | 7a | 0.11 | 0.13 | 0.14 |
| | 3g | 0.12 | 0.13 | 0.06 |
| | 3e | 0.11 | 0.13 | 0.13 |
| **Oman 95 75** | | | | |
| | **Population** | | | |
| | O1 | 0.15 | 0.16 | 0.07 |
| | O2 | 0.16 | 0.15 | -0.04 |
| | O3 | 0.15 | 0.16 | 0.11 |
| | O4 | 0.16 | 0.16 | 0.01 |
| | O5 | 0.19 | 0.16 | -0.16 |
| | **ORF** | | | |
| | 7a | 0.16 | 0.16 | 0 |
| | 3g | 0.18 | 0.15 | -0.18 |
| | 3e | 0.16 | 0.16 | -0.05 |
| **Oman 75 75** | | | | |
| | **Population** | | | |
| | O1 | 0.22 | 0.24 | 0.08 |
| | O2 | 0.19 | 0.23 | 0.15 |
| | O3 | 0.18 | 0.24 | 0.24 |
| | O4 | 0.20 | 0.24 | 0.18 |
| | O5 | 0.23 | 0.24 | 0.05 |
| | **ORF** | | | |
| | 7a | 0.21 | 0.24 | 0.13 |
| | 3g | 0.22 | 0.24 | 0.07 |
| | 3e | 0.21 | 0.24 | 0.13 |
| **Polynesia 95 75** | | | | |
| | **Population** | | | |
| | MH | 0.04 | 0.02 | -0.52 |
| | MT | 0.05 | 0.06 | 0.14 |
| | MV | 0.04 | 0.03 | -0.34 |
| | TF | 0.05 | 0.06 | 0.09 |
| | TV | 0.03 | 0.02 | -0.44 |
| **Polynesia 75 75** | | | | |
| | **Population** | | | |
| | MH | 0.04 | 0.04 | -0.15 |
| | MT | 0.05 | 0.08 | 0.39 |
| | MV | 0.05 | 0.05 | 0.02 |
| | TV | 0.05 | 0.04 | -0.17 |

**Table S5 :** estimate of the index association $\bar{r}_d$ to study linkage disequilibrium in the different datasets. The first column gives the mean and the second the standard deviation computed over 10 000 replicates of 320 SNPs. The "global" results correspond to analyses done at the level of the whole corresponding dataset. The "strata" results correspond to analyses done at the level of mitochondrial lineages for the All and Oman datasets, and at the level of sampling site for the Polynesia datasets (i.e. the "strata" levels used in the poppr R package). Note that the analysis was not done for 5a in Oman where only one individual was analysis.

| **Dataset** | **mean** | **s.d.** |
|---|---|---|
| **Global** | | |
| All_95_75 | 0.033 | 0.005 |
| All_75_75 | 0.099 | 0.019 |
| Oman_95_75 | 0.002 | 0.001 |
| Oman_75_75 | 0.004 | 0.001 |
| Polynesia_95_75 | 0.035 | 0.007 |
| Polynesia_75_75 | 0.017 | 0.007 |
| **Strata All_95_75** | | |
| 3e | 0.003 | 0.006 |
| 3g | 0.025 | 0.025 |
| 5a | 0.019 | 0.004 |
| 7a | 0.005 | 0.002 |
| **Strata All_75_75** | | |
| 3e | 0 | 0.003 |
| 3g | 0.049 | 0.009 |
| 5a | 0.083 | 0.017 |
| 7a | 0.007 | 0.002 |
| **Strata Oman_95_75** | | |
| 3e | 0.003 | 0.003 |
| 3g | 0.002 | 0.006 |
| 7a | 0.003 | 0.001 |
| **Strata Oman_75_75** | | |
| 3e | 0 | 0.002 |
| 3g | 0.05 | 0.006 |
| 7a | 0.007 | 0.001 |
| **Strata Polyn_95_75** | | |
| MH | 0.278 | 0.154 |
| MT | 0.008 | 0.009 |
| MV | 0.014 | 0.024 |
| TF | 0.086 | 0.043 |
| TV | 0.116 | 0.101 |
| **Strata Polyn_75_75** | | |
| MH | 0.235 | 0.072 |
| MT | 0.01 | 0.011 |
| MV | 0.008 | 0.014 |
| TV | 0.108 | 0.065 |

**Figure S1** : histograms of genetic distance among individuals for the different datasets. The distances correspond to the proportion of differences among loci computed with poppr. We indicate below each histogram the retained threshold to remove repeated MLLs.

**A) All_95_75**

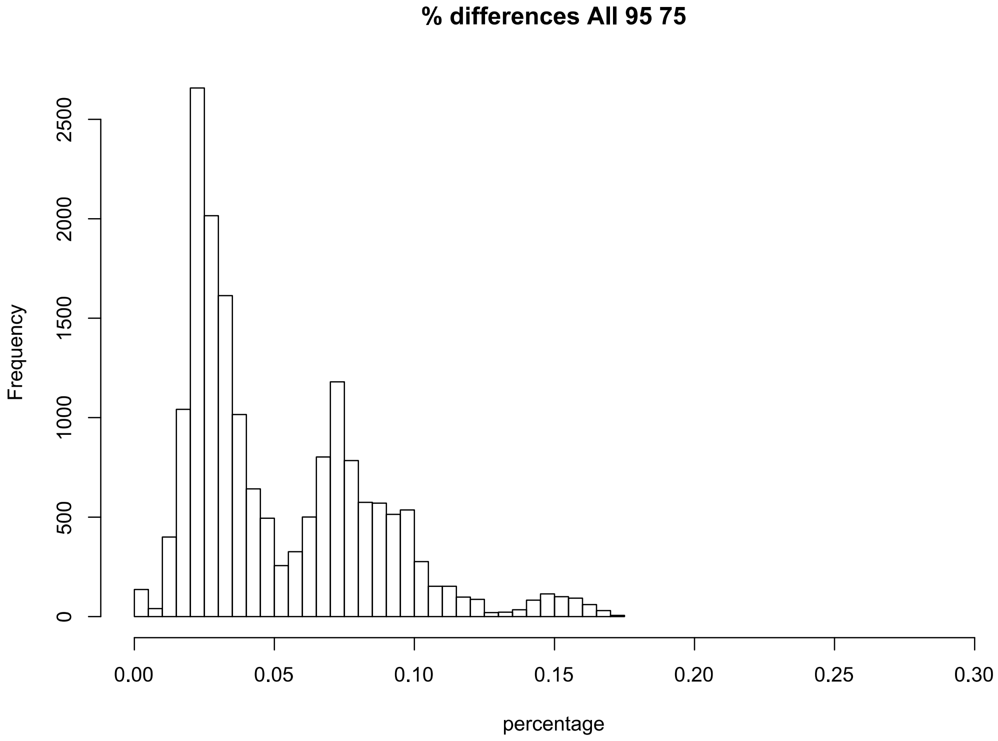


MLL threshold 0.0157

**B) All_75_75**

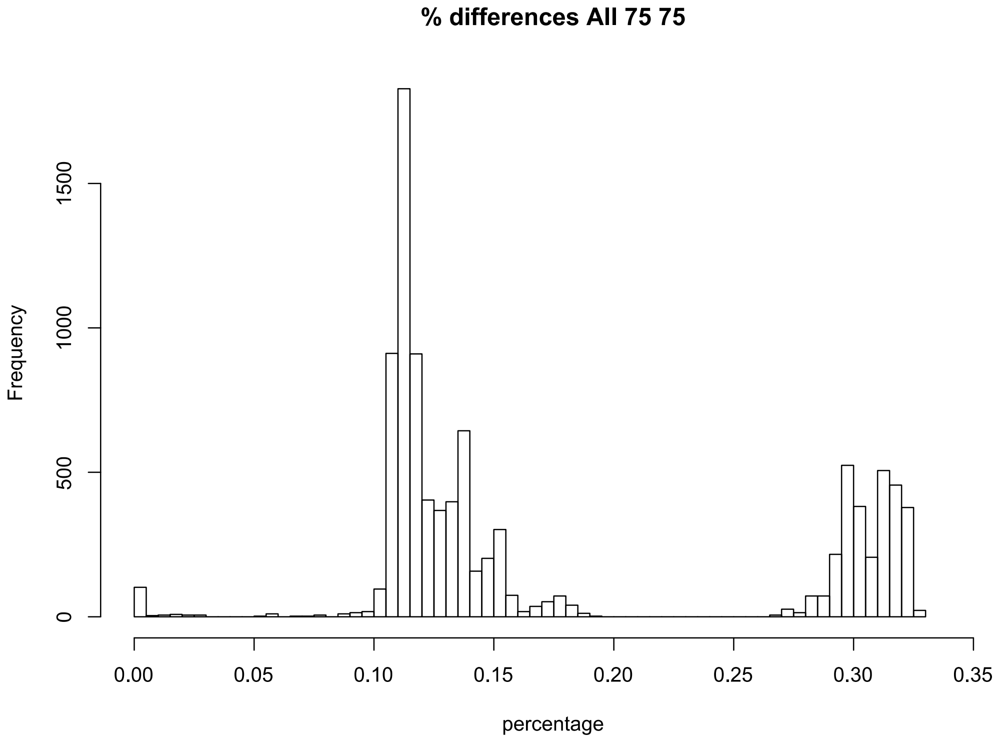


MLL threshold 0.06

**C) Oman_95_75**

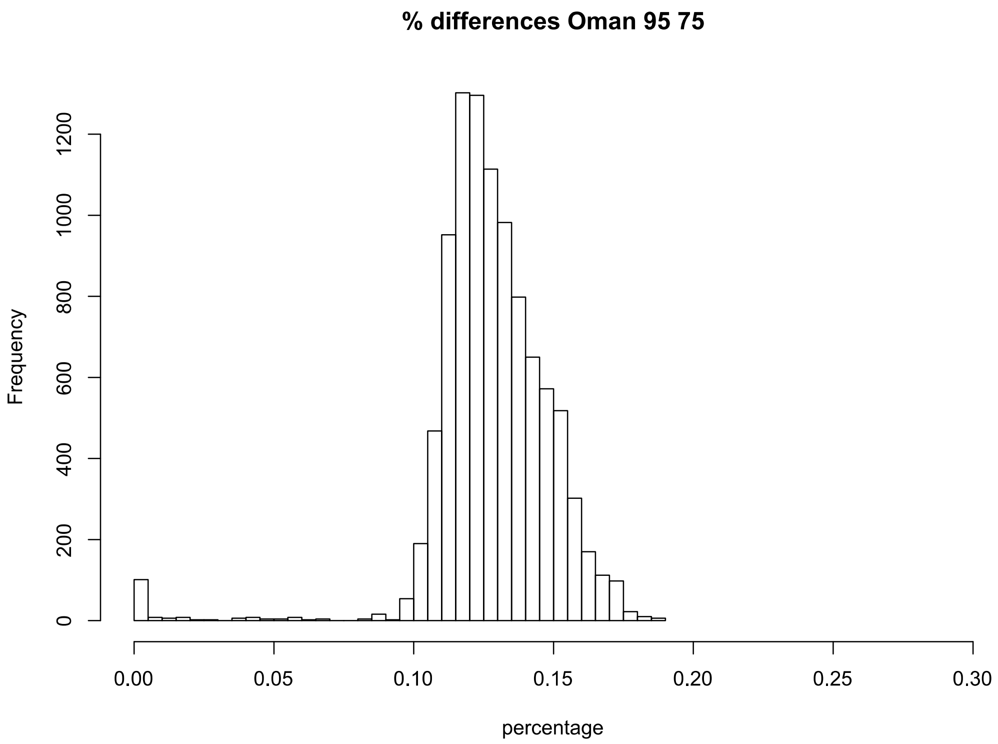


MLL threshold 0.0685

**D) Oman_75_75**

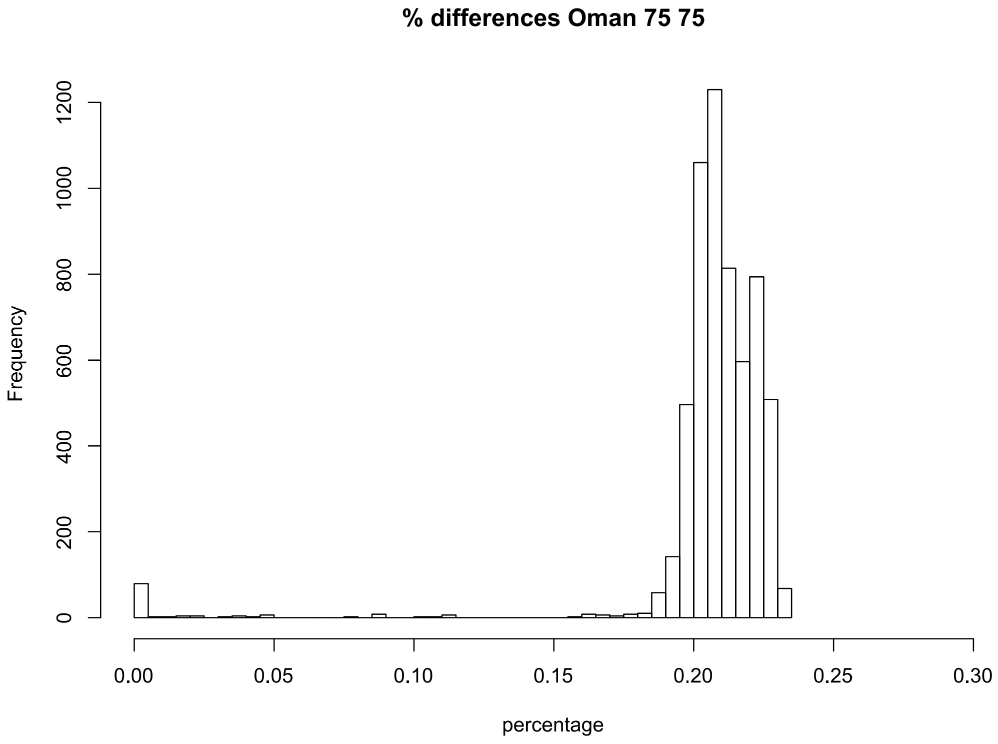


MLL threshold 0.09

**E) Polynesia_95_75**

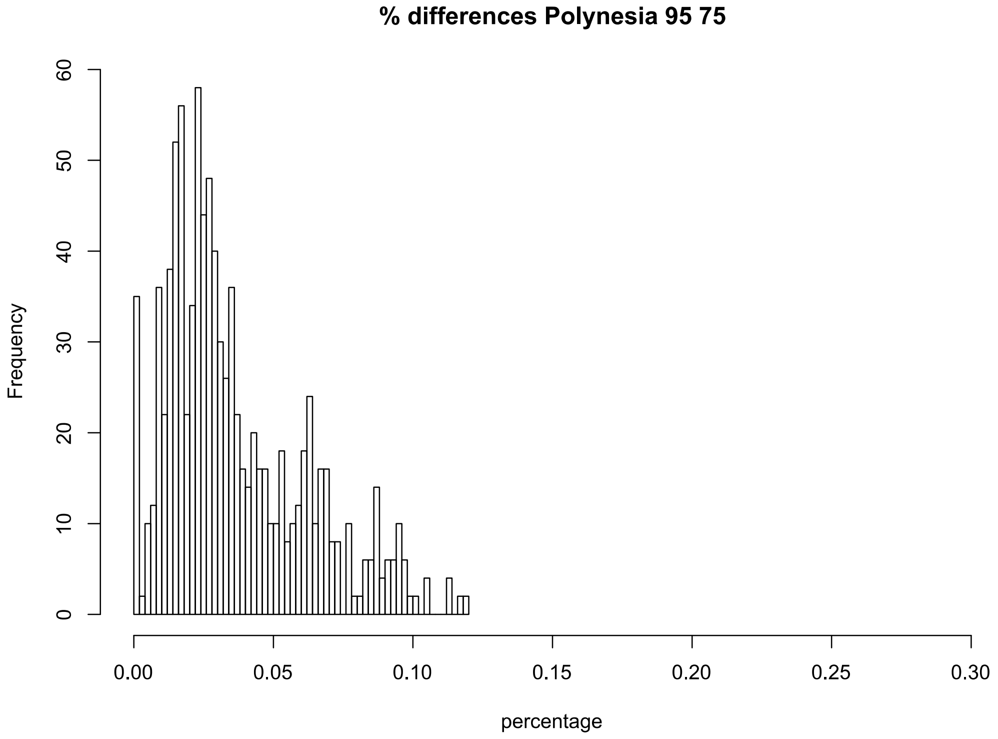


MLL threshold 0.002

**F) Polynesia_75_75**

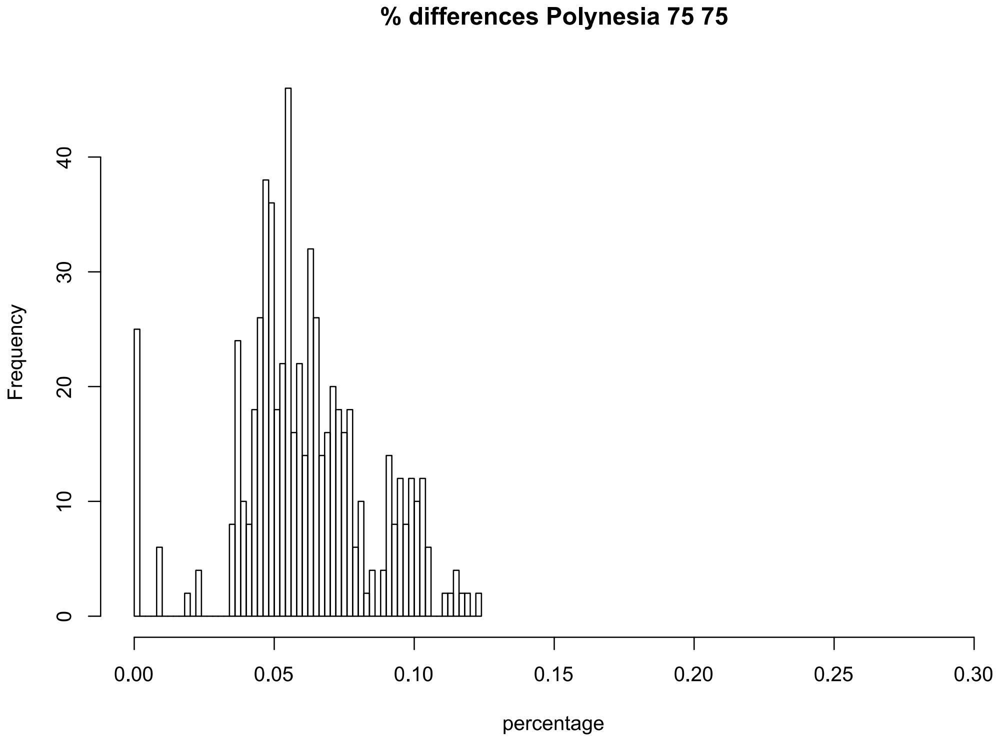


MLL threshold 0.04

**Figure S2:** networks based on the percentage of difference among individuals for the All, Oman and Polynesia 75_75 datasets. The colors indicate the corresponding species hypothesis according to the sequence of mitochondrial ORF.

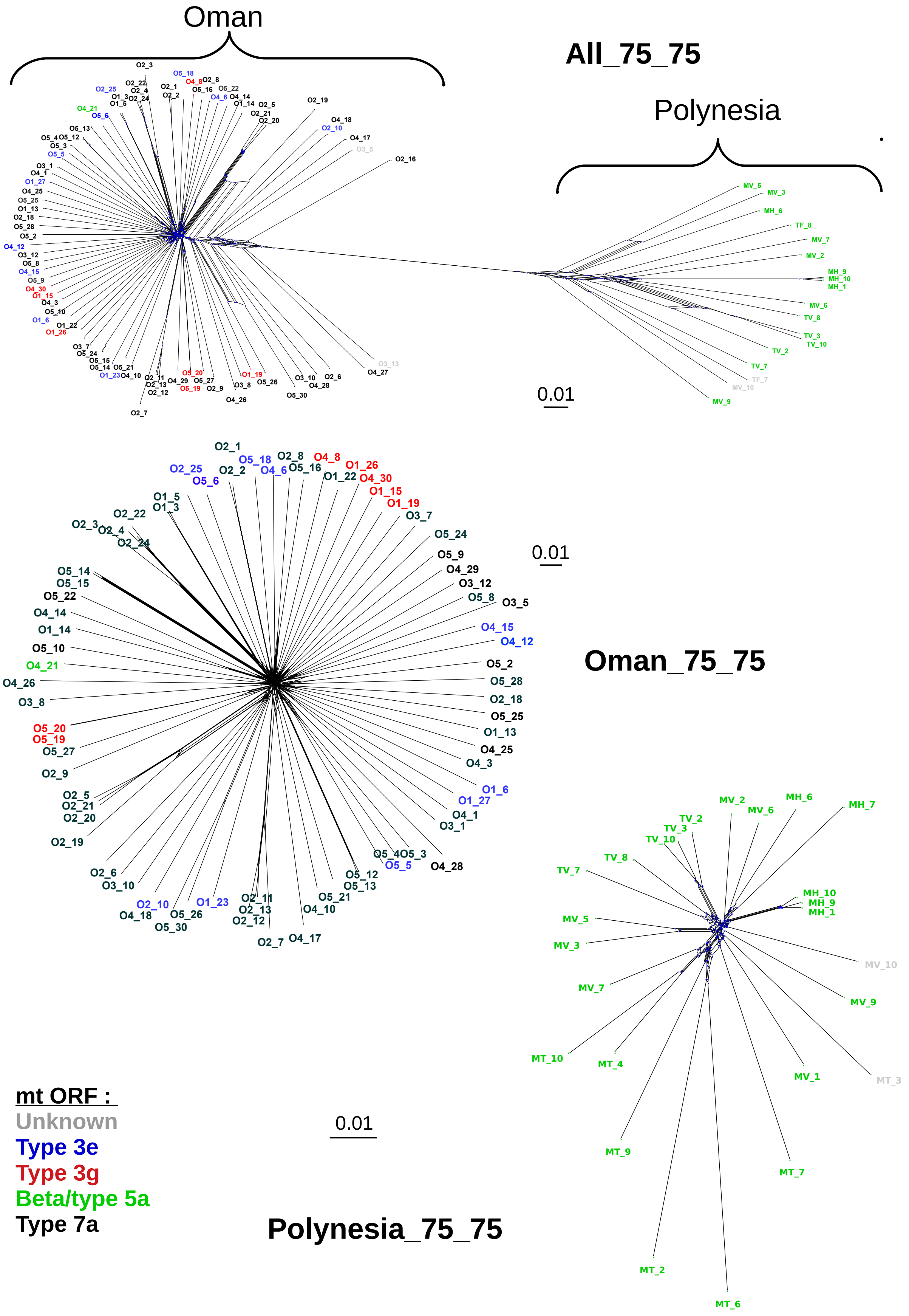

**Figure S3** : histograms of individual inbreeding coefficients F estimated with VCFtools for the different datasets. The blue histogram corresponds to all individuals, and the yellow histograms corresponds to individuals potentially involved in MLLs with the previously defined distance thresholds. For visibility the x axis is different among figures. A red line at x = 0 is given for comparison.

**All_95_75**

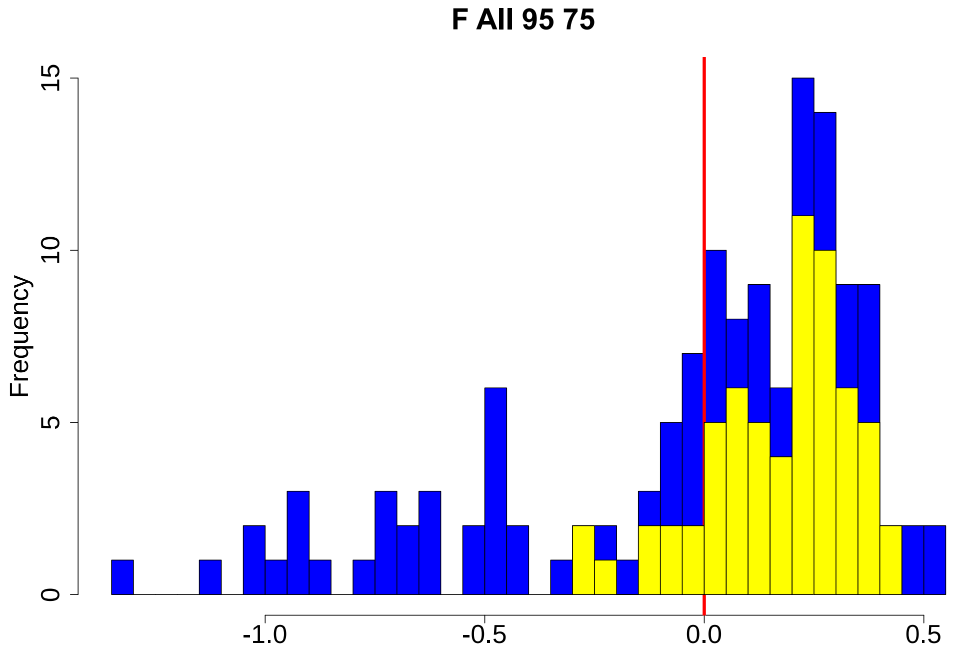


**All_75_75**

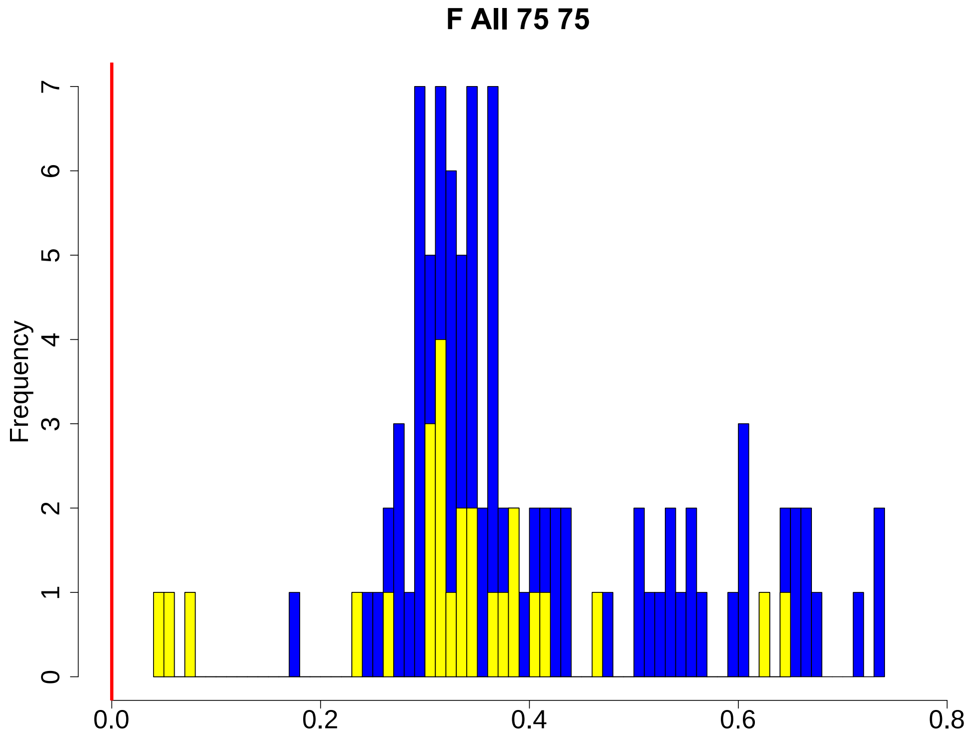

**Oman_95_75**

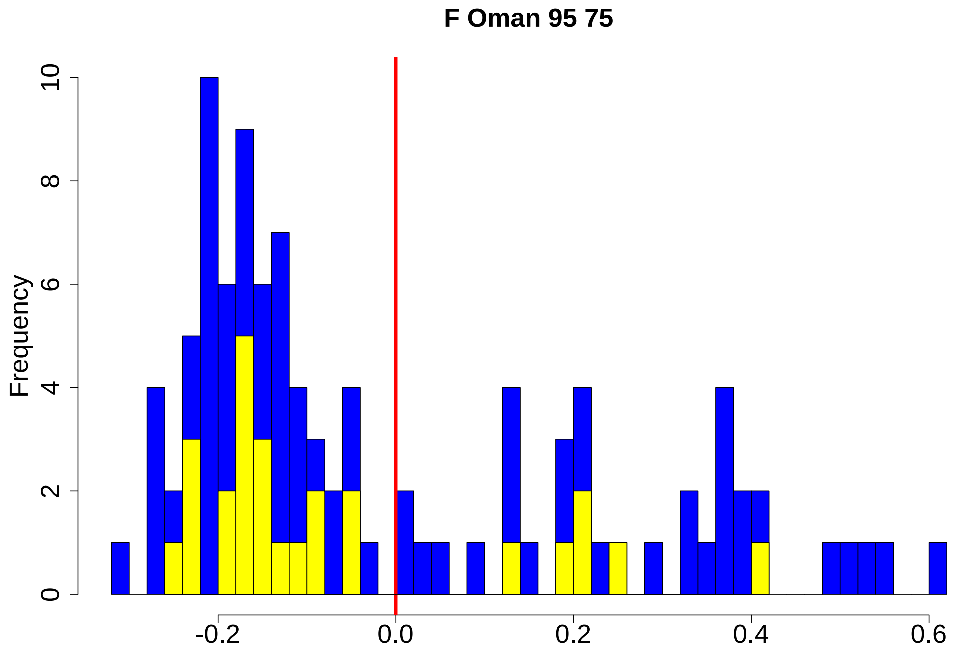


**Oman_75_75**

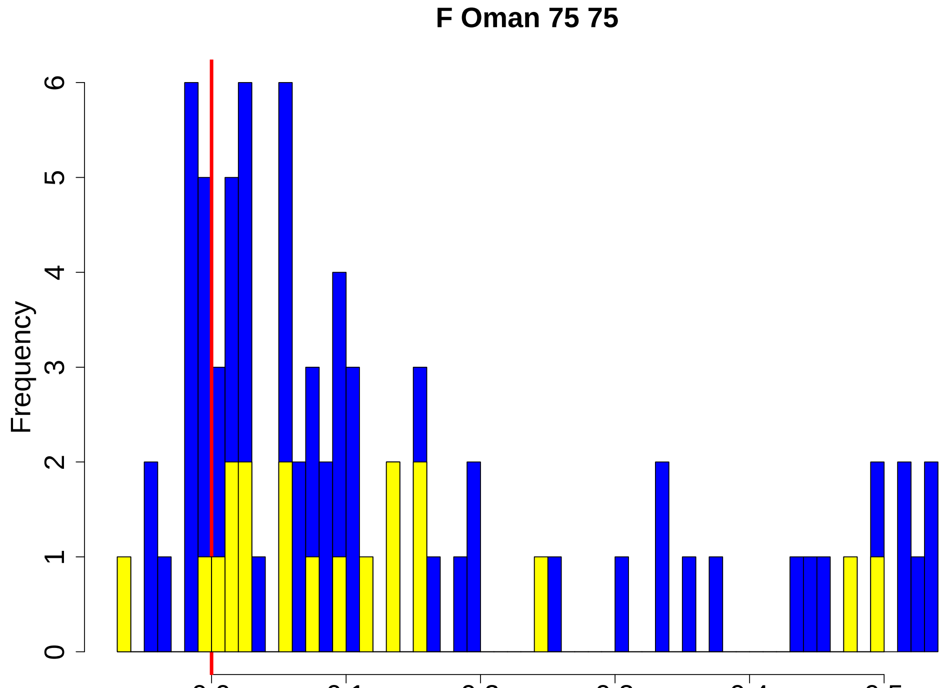

**Polynesia_95_75**

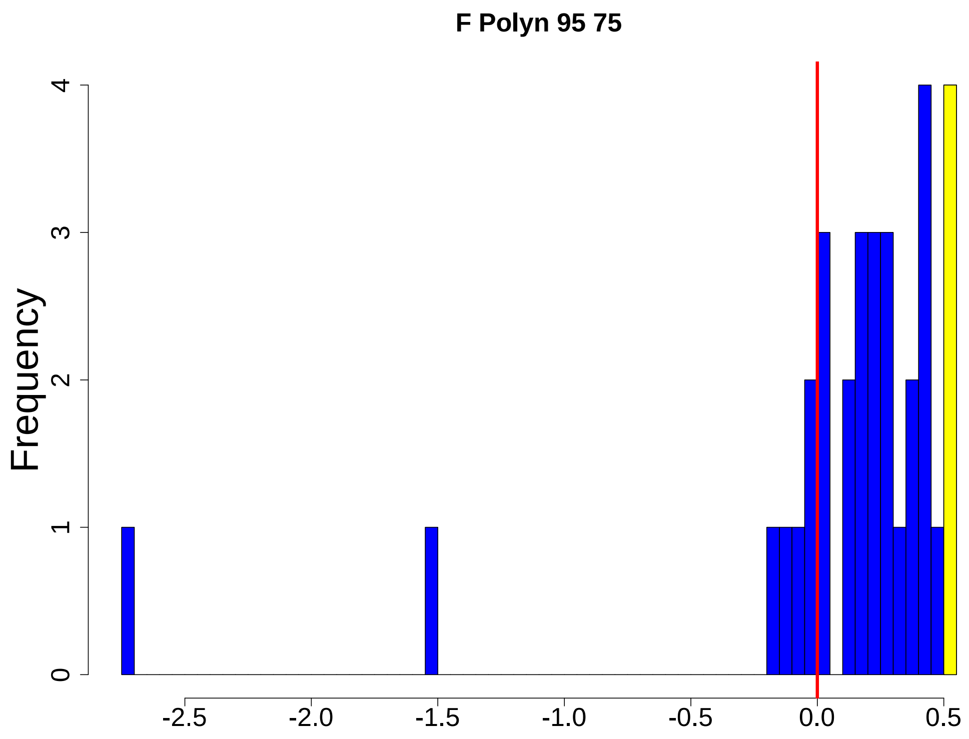


**Polynesia_75_75**

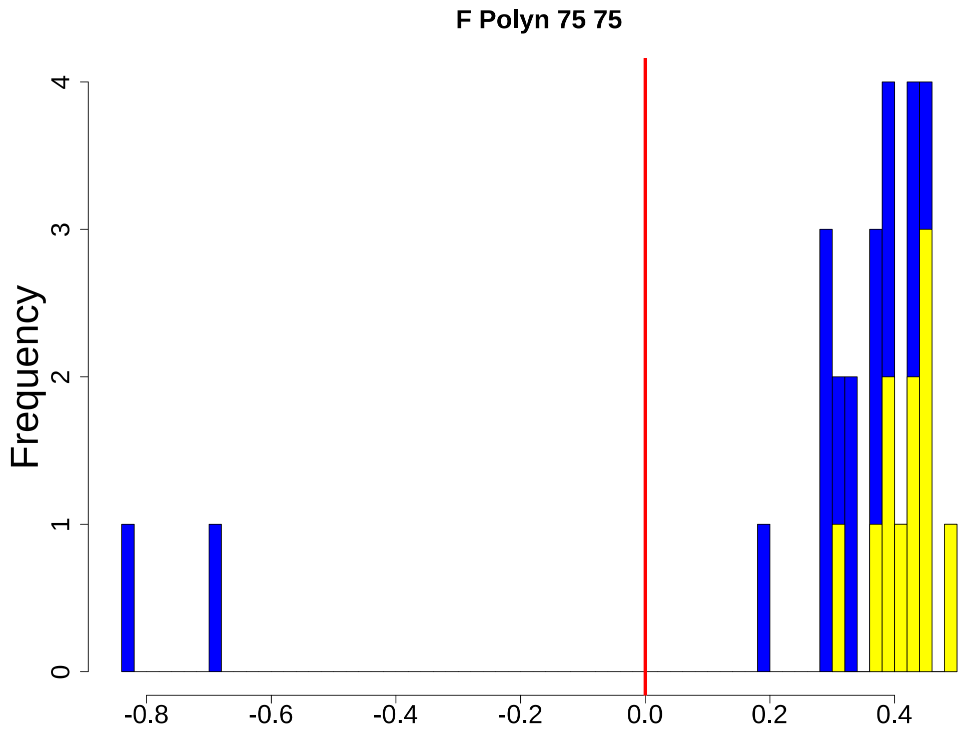

**Figure S4 :** distribution of the estimates of the index association $\bar{r}_d$ to study linkage disequilibrium in the different datasets : A) on the whole datasets, B) per mitochondrial lineages or per site. See the main text and the legend of Table S4 for details.

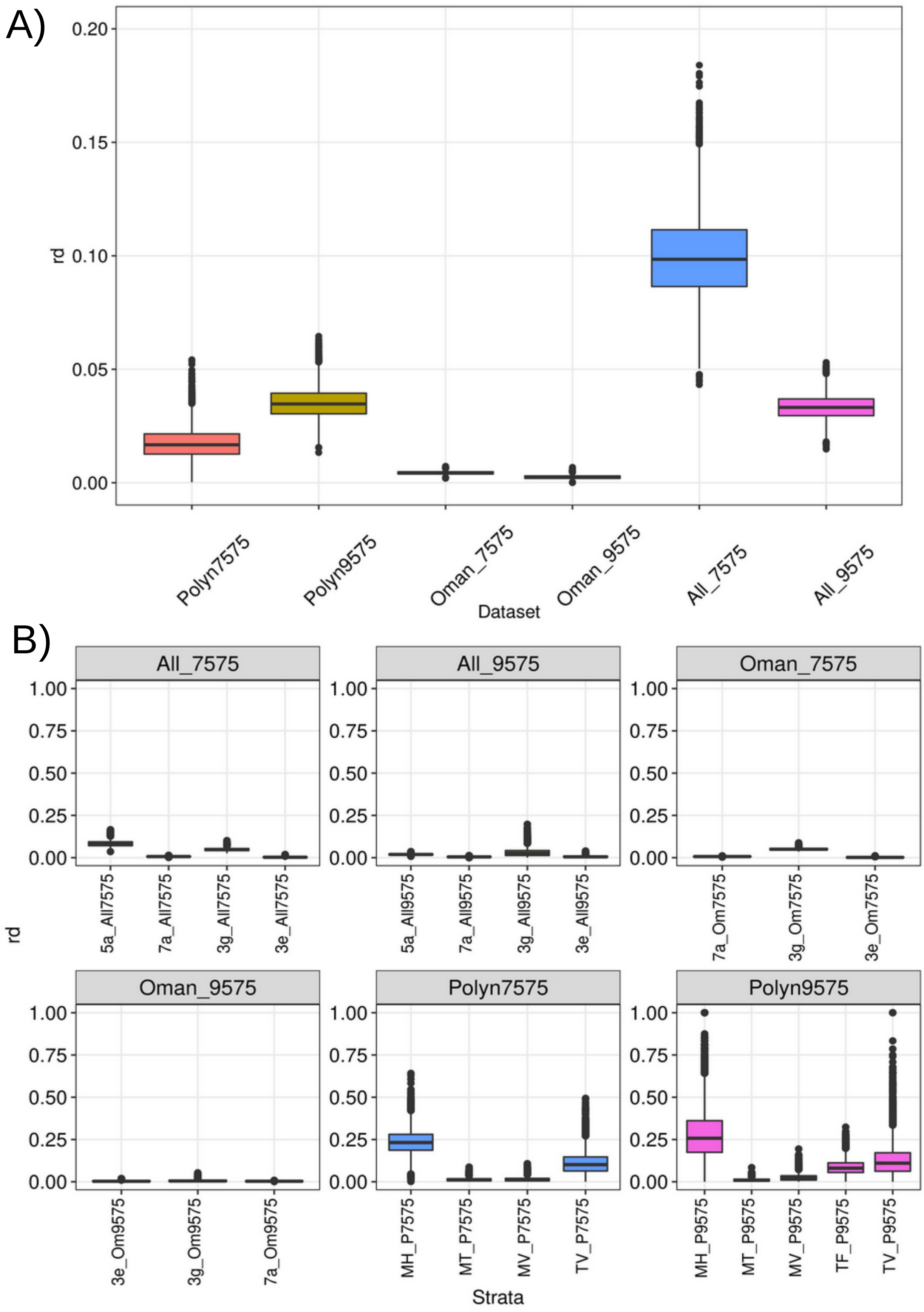

**Figure S5:** distribution of $F_{ST}$ over all loci for the comparison between Oman and Polynesia for A) the All_95_75 dataset (mean $F_{ST}$ estimate: 0.105) and B) the All_75_75 dataset (mean $F_{ST}$ estimate: 0.352).

**A)**

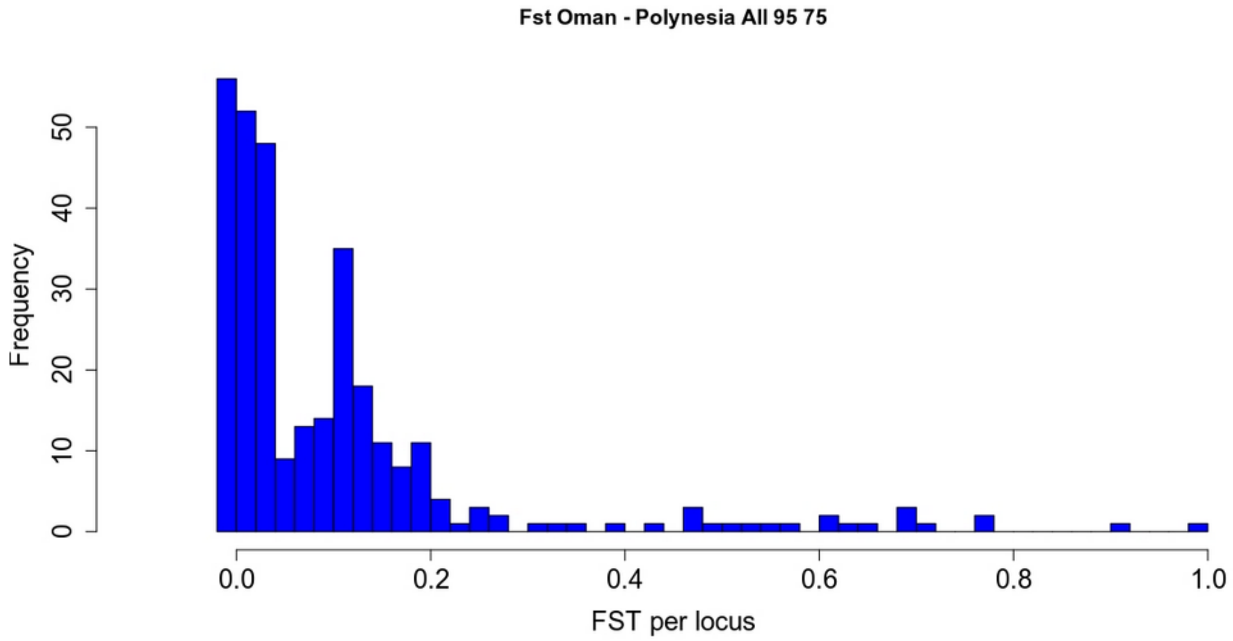


**B)**

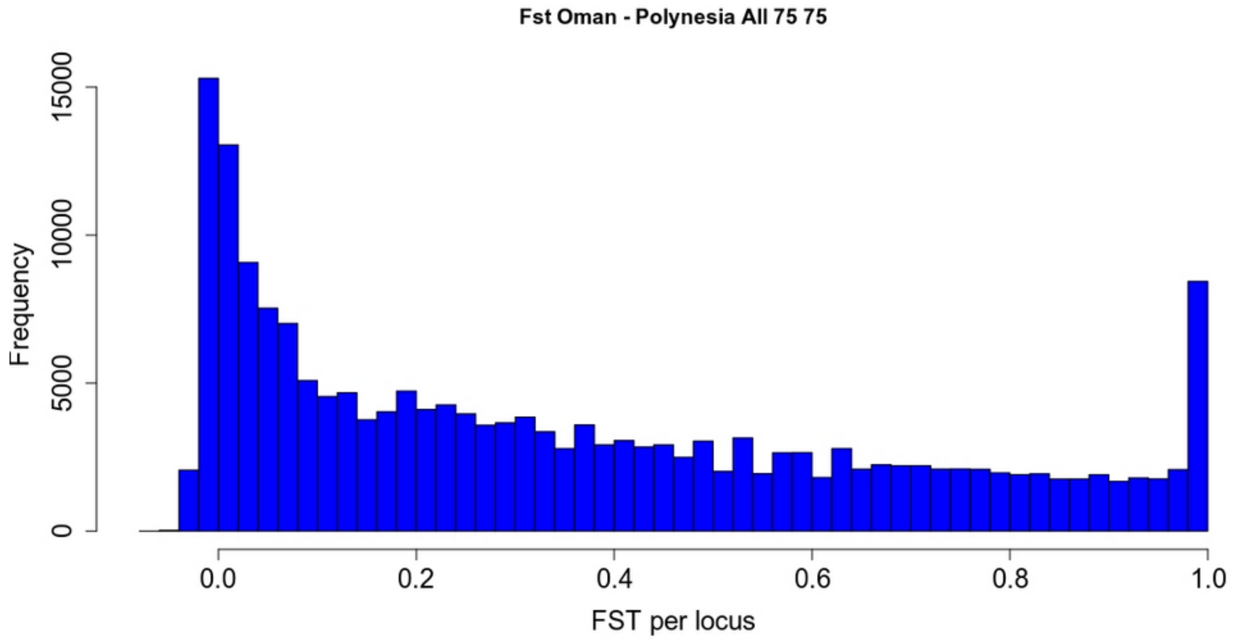

**Figure S6:** plot of individuals on the PCA axes 1 and 2 for A) the All_95_75, and B) the All_75_75 datasets. For All_95_75, the percentage of inertia was 9.2 for axis 1 and 5.6 for axis 2. For All_75_75, the percentage of inertia was 25.1 for axis 1 and 3.9 for axis 2. The individual dots are colored according to their ORF mitochondrial lineage, and a focus on the central part is given under each plot.

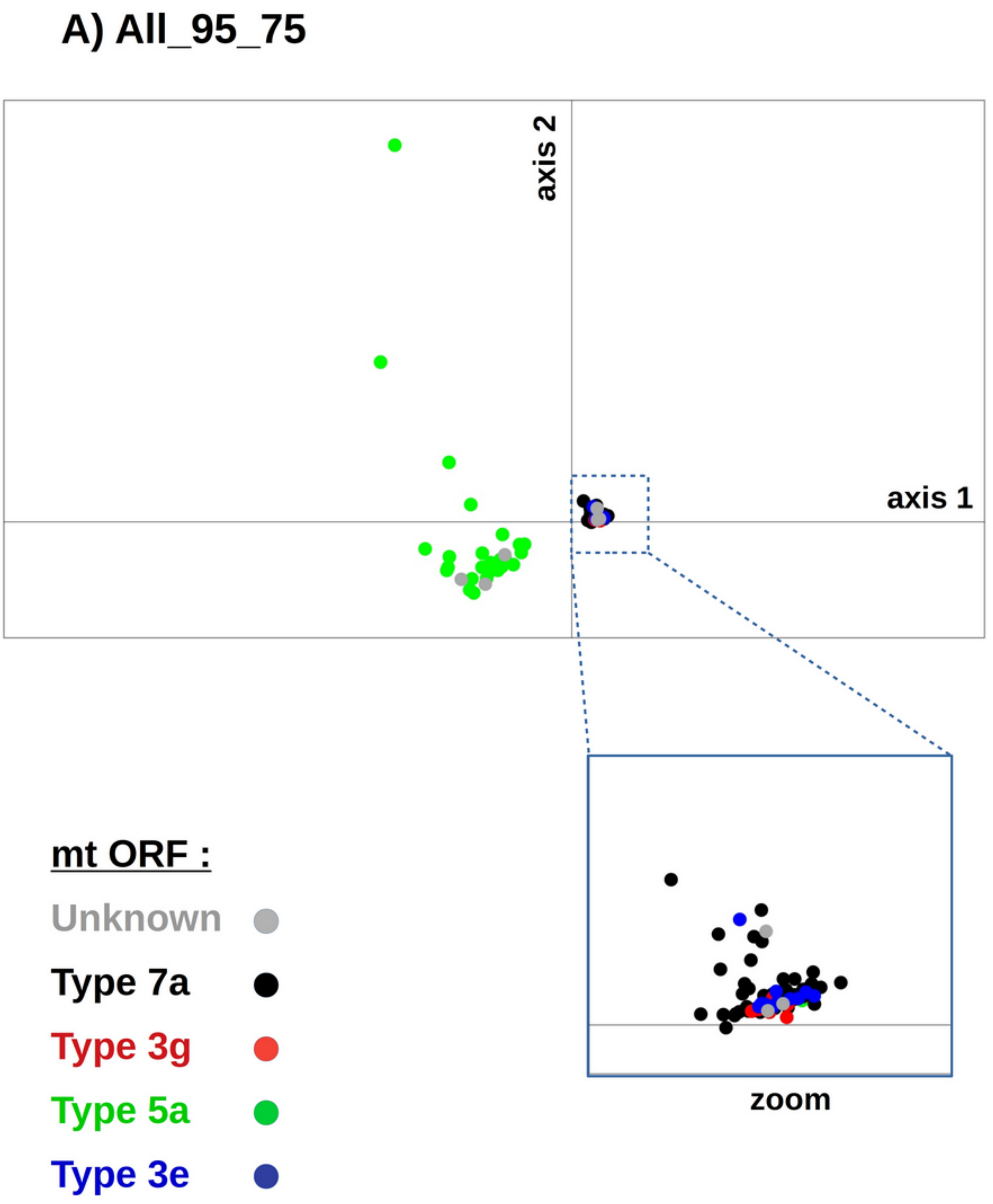

### B) All_75_75

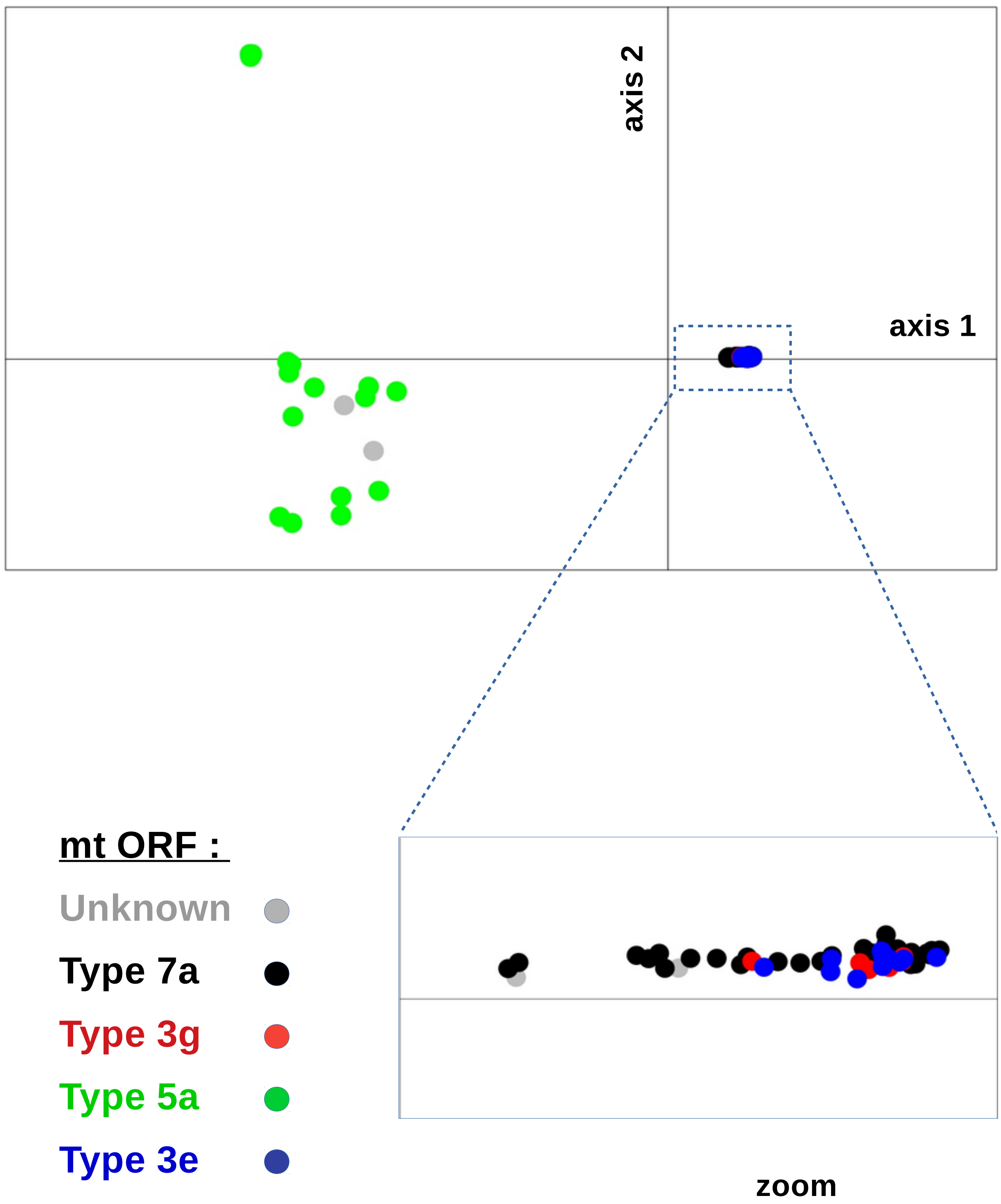

axis 2
axis 1
mt ORF :
Unknown
Type 7a
Type 3g
Type 5a
Type 3e
zoom

**<u>Simulations : distribution of *F* estimates :</u>**

The figures below show the distribution of the average (A), maximum (B) and minimum (C) of the *F* parameter over individuals for different simulation configurations, with 30 simulations each. These distributions are compared to observed average, maximum and minimum values displayed as horizontal lines for the different datasets (see legends). Notation of the simulation configurations : panm : panmixia ; clon01, clon05, and clon09 : clonality present at rates 0.1, 0.5 and 0.9 respectively ; self_clon01 and self_clon05 : selfing present at rate 0.1, and clonality present at rates 0.1 and 0.5 respectively.

**A)** average of *F* parameter over simulation replicates ; horizontal lines give observed values for the different datasets : green : all, red : Oman, blue : Polynesia, continuous line : 75_75, dotted line : 95_75.

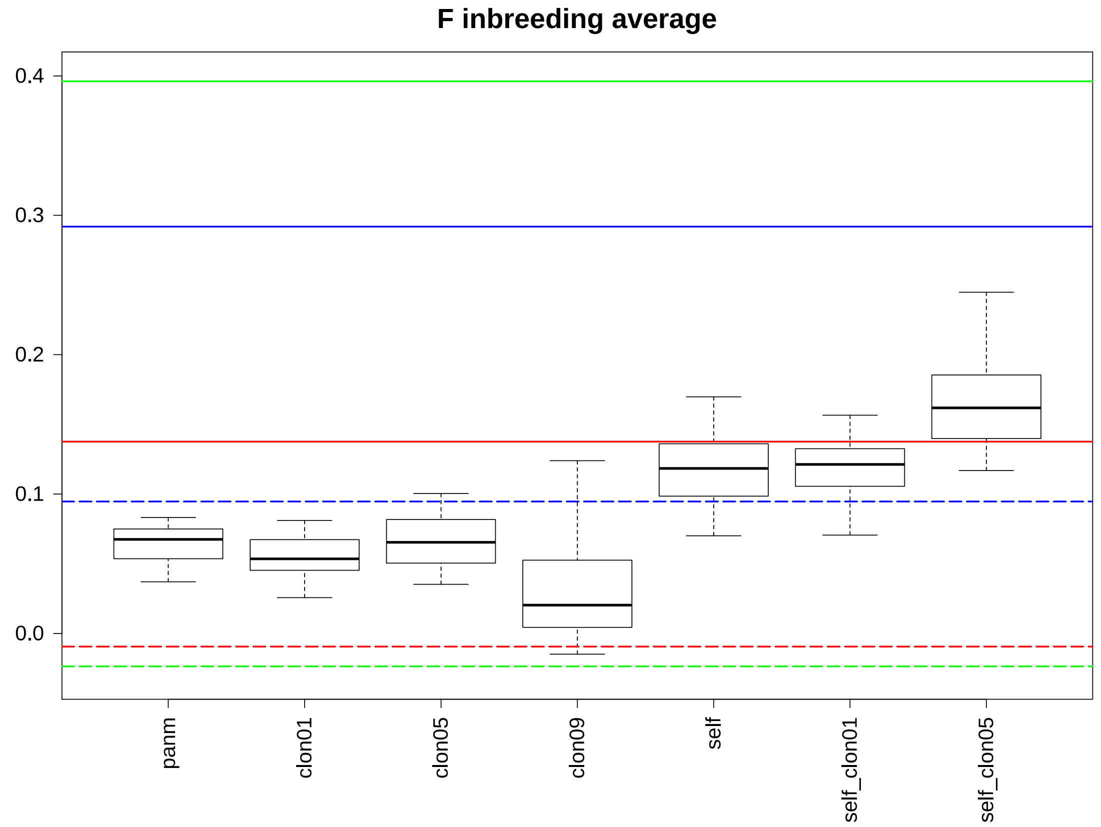

**B)** maximum of *F* parameter over simulation replicates ; horizontal lines give observed values for the different datasets : green : all, red : Oman, blue : Polynesia, continuous line : 75_75, dotted line : 95_75.

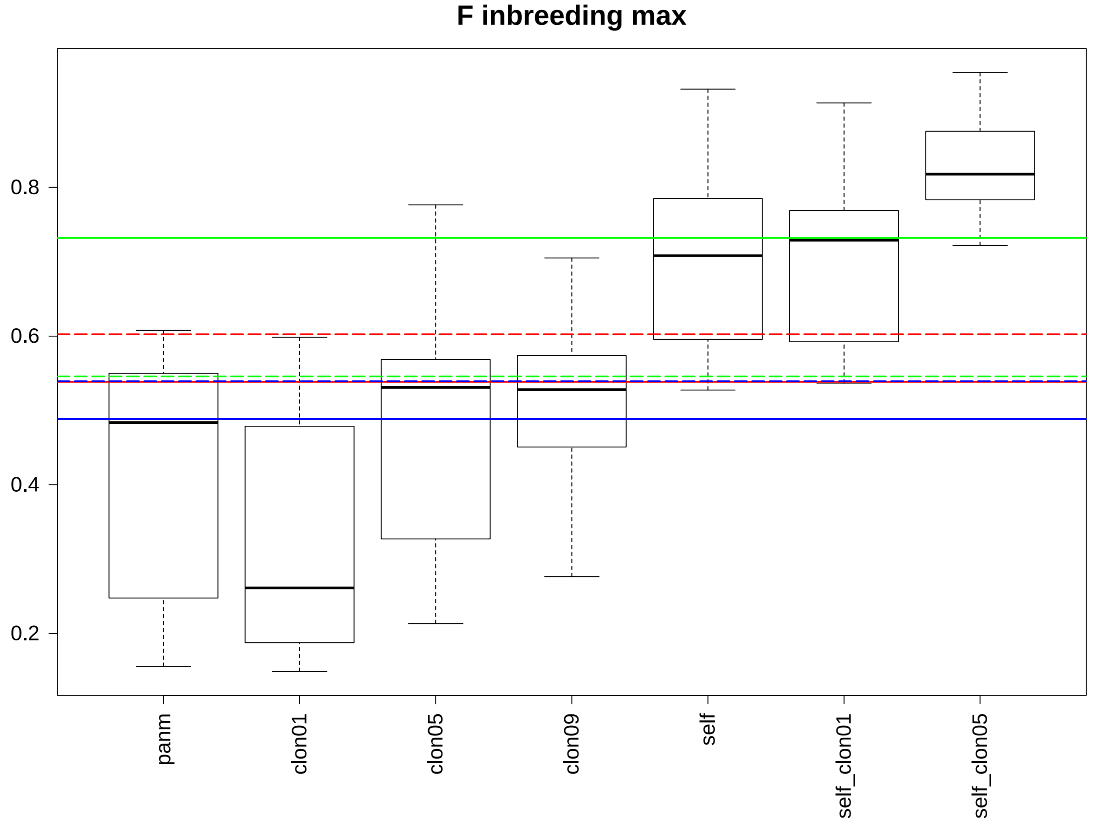

**C)** minimum of *F* parameter over simulation replicates ; horizontal lines give observed values for the different datasets : green : all ; red : Oman ; blue : Polynesia ; continuous line : 75_75 ; dotted line : 95_75.

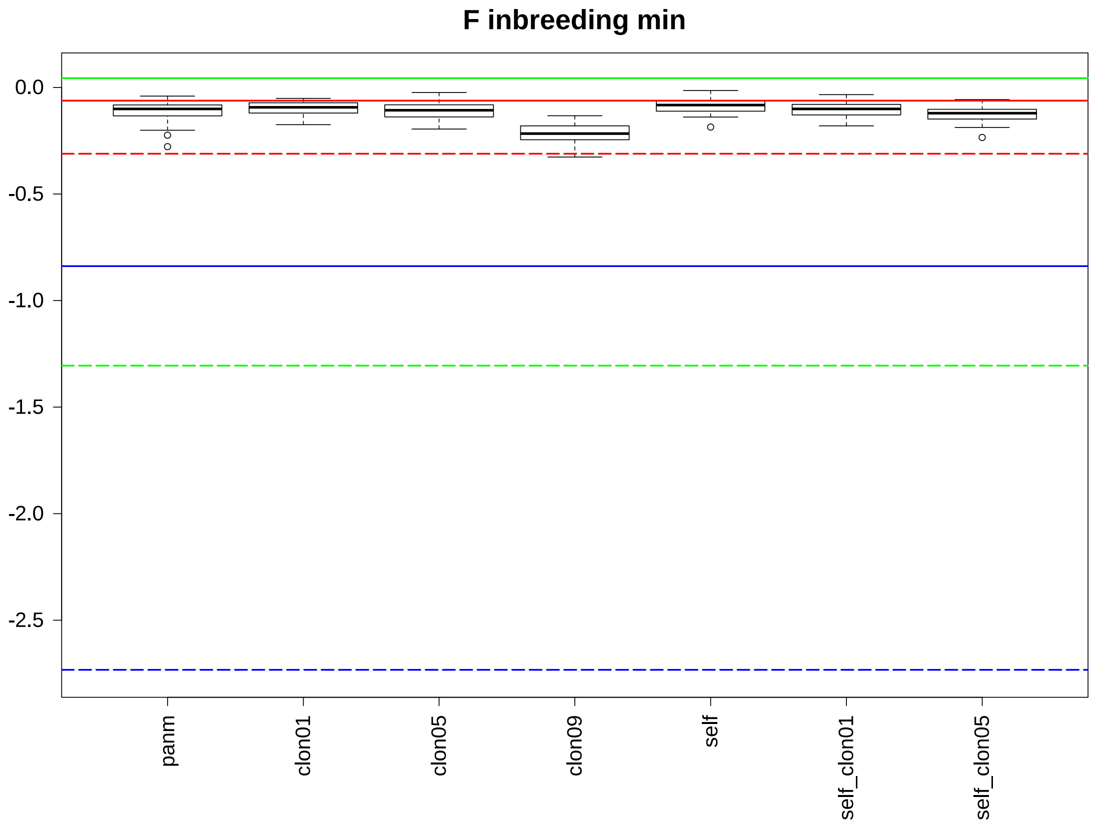

**Simulations : analysis of linkage disequilibrium :**

The figures below show the distribution of the mean (A) and standard deviation (B) of $\bar{r}_d$ over simulations. Note that $\bar{r}_d$ was here computed separately for the two simulated populations in each simulation,with mean and standard deviations computed across subsampling replicates in each population. These distribution are compared to samples with the two highest observed mean values obtained in the 75_75 datasets at the level of populations or mitochondrial lineages : All_75_75 : 3g (light green) and 5a (dark green) ; Oman_75_75 : 3g (red) and 7a (orange) ; Polynesia_75_75 : MH (light blue) and TV (dark blue) ; close values

**A) mean of** $\bar{r}_d$ ; the lines for All_75_75 3g (light green) and Oman_75_75 3g overlap (red).

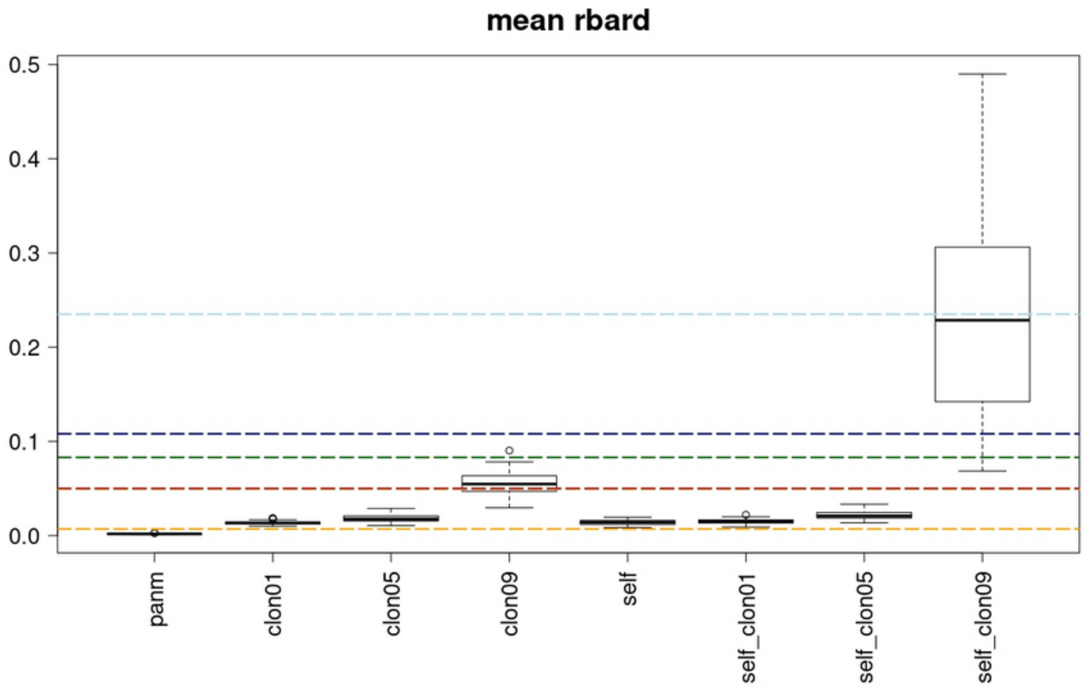

**B)** standard deviation of $\bar{r}_d$ ; for clarity we did not plot the lines for Polynesia_75_75 MH (s.d. = 0.072) and TV (s.d. = 0.065).

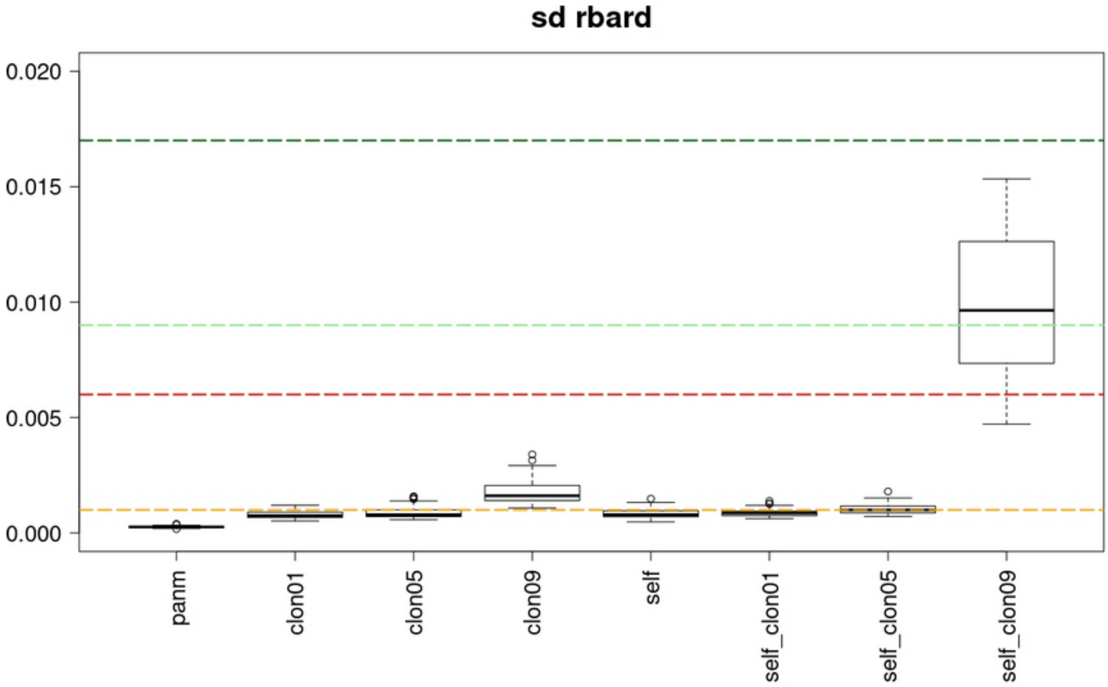

**Simulations : distribution of $F_{ST}$ estimates :**

The figures below show the distribution of the $F_{ST}$ (average over loci) between the two populations over 30 simulations, for the different simulation configurations. These distribution are compared to observed average, maximum and minimum values displayed as horizontal lines for the different datasets : green : all, red : Oman, blue : Polynesia, continuous line : 75_75, dotted line : 95_75.

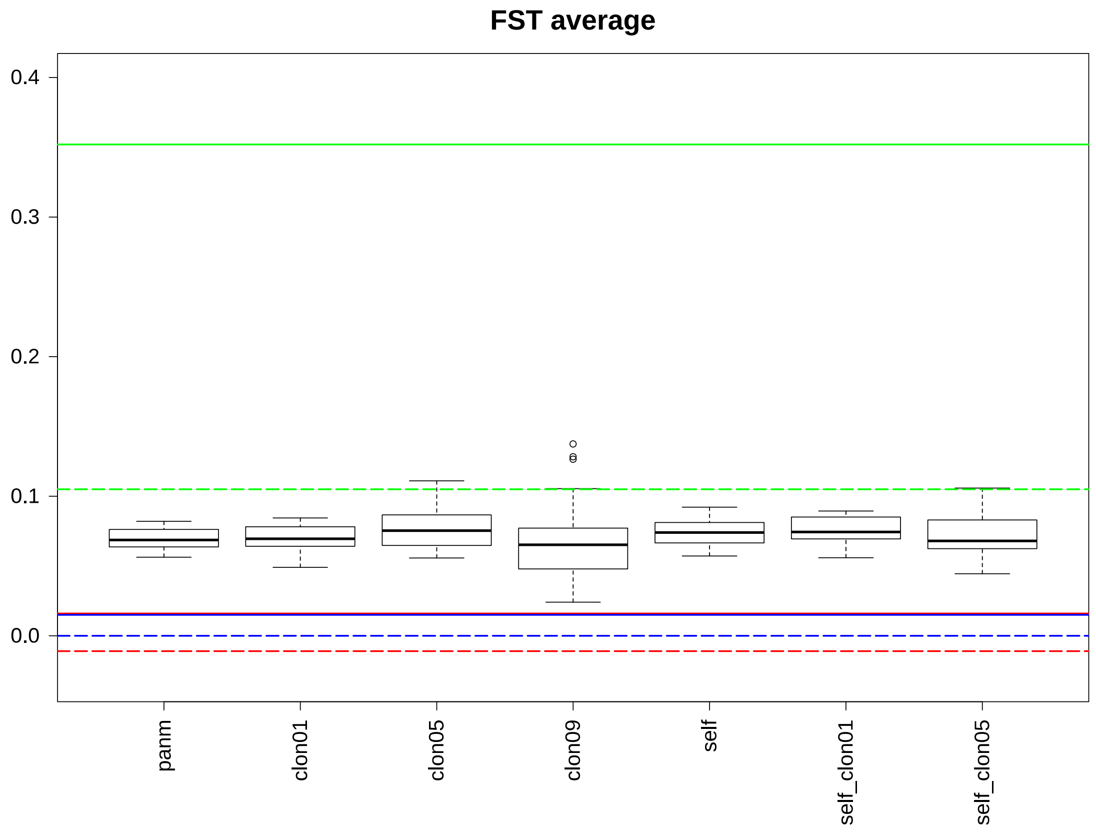
